# Periventricular and Deep White Matter Hyperintensity Thresholds in Aging: Exponential Progression, Cognitive Decline, and Neuroanatomic Atrophy

**DOI:** 10.1101/2024.09.03.610993

**Authors:** Niraj Kumar Gupta, Neha Yadav, Vivek Tiwari

**Affiliations:** Indian Institute of Science Education and Research (IISER) Berhampur, India

**Keywords:** Aging, Cerebral Small Vessel Disease, White Matter Hyperintensity, Cognitive Impairment, Neuroanatomic Atrophy, Cognitive performance

## Abstract

White matter hyperintensities (WMH), which are brain lesions associated with cerebral small vessel disease and aging, signify fiber loss and pruning. Analysis of T2-FLAIR MRI data from the NACC cohort, including cognitively normal (CN), cognitively impaired (CI), and Alzheimer’s disease (CI-AD) subjects, revealed that a significant subset of participants, even those classified as CN, harbor substantial periventricular (PVWMH) and deep white matter hyperintensity (DWMH) loads, while others displayed minimal or no PVWMH and DWMH, across ages 50-94 years. In this study, we quantified the thresholds and progression kinetics of PVWMH and DWMH and their impact on cognitive performance and neuroanatomic changes in the aging cohort (NCN = 521, NCI = 146, NCI-AD = 319). Our findings explore the impact of PVWMH and DWMH loads on global and specific cognitive domains to determine whether cognitive impairments are directly induced by PVWMH and DWMH loads, or mediated through distinct neuroanatomic structures. PV and DWMH loads are higher in CI and CI-AD subjects compared to CN but the PVWMH and DWMH loads are not discriminative of CI and CI-AD. The progression kinetics of PVWMH and DWMH volume with age indicate an exponential rate of increase, with PVWMH escalating approximately twice as fast as DWMH particularly around an inflection point at 61 years of age. PVWMH load presents with increased probability of occurrence in frontal horn compared to occipital horn while DWMH is diffused and accumulates significantly at later age than that observed for PVWMH. Multivariate global regression suggested significant effect (p<0.01) of PVWMH on Trail making tests (TMTs)-A and B (executive function), animal naming tests (semantic memory), with no significant effects observed for DWMH load. Indeed, beyond a threshold of PVWMH volume >2.3 mL, significant deficits in TMTs were observed compared to the subjects without PVWMH load. A PVWMH volume >2.3 mL and DWMH >2.75 mL is significantly associated with impairments in attention & working memory (Digit Span Tests), and semantic memory. Noticeably, significant neuroanatomic atrophy in the cerebral cortex, nucleus accumbens, RMFG, precentral, and paracentral gyrus is observed for PVWMH load >2.3 mL, while DWMH load was not significantly associated with neuroanatomic loss. Furthermore, a mediation model employing neuroanatomic volumes as mediator, PVWMH load as predictor and cognitive tests as outcome suggested that PVWMH volume contributes significantly to deficits in TMT-B mediated through atrophy in precentral gyrus (64%), accumbens (39%), paracentral gyrus (32%), rostral middle frontal gyrus (31%), and lingual gyrus (30%), each contributing distinct proportions, alongside the direct effect. DWMH load did not emerge as a significant predictor (direct or indirect) for the cognitive deficits. Further, no significant neuroanatomic mediations from PVWMH load were observed for other cognitive tests indicative of direct involvement of PVWMH load. Global cognition like MMSE was affected only at a higher PVWMH accumulation (>6 ml).

**Highlights:**

- PVWMH escalates exponentially twice as fast as DWMH with age.
- PVWMH >2.3 mL is linked to cognitive deficits in executive function, semantic memory and neuroanatomic atrophy; DWMH less impactful.
- PVWMH, not DWMH, significantly affects cognitive decline via a unique set of brain structural loss.
- PVWMH and DWMH loads are not discriminative of CI and AD subjects.

**Overview:** 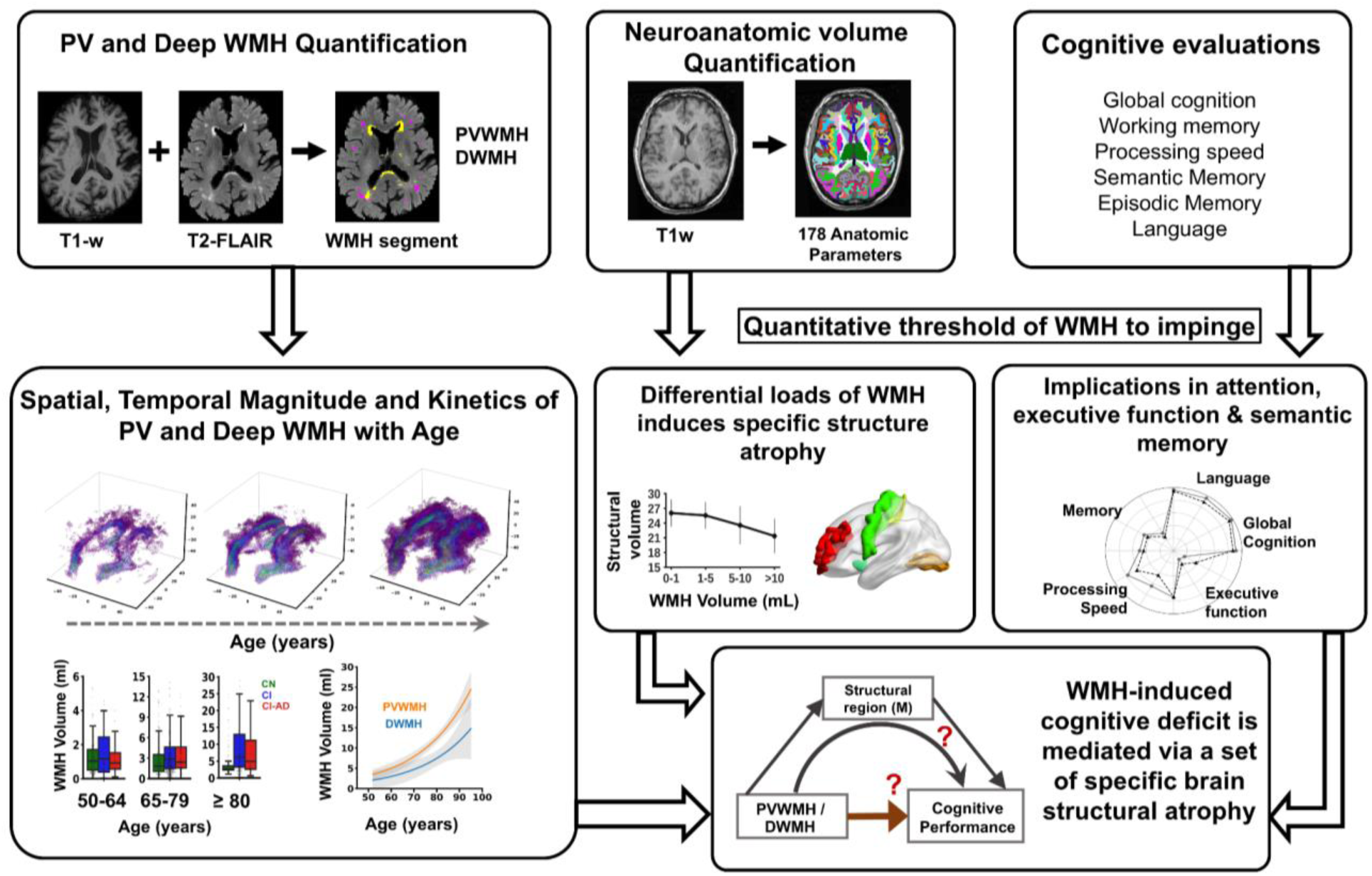

## Introduction

Cerebral Small Vessel Disease (CSVD) is an aging-related brain pathology, arising from small vessel infarcts, leading to fiber pruning and loss (Pantoni, 2010; Wardlaw et al., 2015). The fiber derangement is observed as White Matter Hyperintensities (WMH) on T2-FLAIR MR images (Wardlaw et al., 2019, 2013). The prevalence and volume of WMH increase markedly with advancing age, though there is considerable variability among individuals, wherein a subset of subjects presents with minimal or no WMH in the brain, while another subset of the same age groups presents with substantial WMH load. The conventional WMH assessments, like Fazekas grading (Fazekas et al., 2002, 1987), qualitatively link WMH to neuroanatomic and cognitive risks (Debette and Markus, 2010; Prins and Scheltens, 2015) with aging. Periventricular (PVWMH) and Deep White Matter Hyperintensities (DWMH) reflect ischemic changes detectable through neuroimaging. However, there remains a need for a quantitative approach to define WMH thresholds beyond which there is enhanced vascular insult leading to structural and cognitive alterations. Accurate quantification of PVWMH and DWMH is crucial for establishing a clear risk profile for brain health alterations beyond normal aging.

To investigate the relationship between WMH load and its impact on structural and cognitive domains, we quantified PVWMH and DWMH alongside comprehensive neuroanatomic segmentations and cognitive assessments in cognitively normal (CN), cognitively impaired (CI), and cognitively impaired due to Alzheimer’s disease (CI-AD) subjects across early (50-64 years), intermediate (65-79 years), and late (≥80 years) age groups from the National Alzheimer’s Coordinating Centre (NACC) (Beekly et al., 2007, 2004; Besser et al., 2018) cohort. We have attempted to address a central question whether PVWMH and DWMH load have unique threshold and kinetics with aging and their contributions to perturbations across cognitive domains, global cognition and neuroanatomical health.

Comprehensive segmentation and quantification of PVWMH and DWMH, along with neuroanatomic volumetry, revealed significant insights. PV and DWMH segmentation across CN, CI and CI-AD subjects revealed that even within subjects classified as CN, a substantial percentage of subjects presented with PVWMH and WMH load at all the age groups; early, intermediate and late. The probability of PVWMH and DWMH occurrence increases exponentially with age, with PVWMH escalating approximately twice as fast as DWMH. A threshold of PVWMH load >2.3 ml in cognitively normal subjects is implicated in impaired cognition associated with attention (poor performance in Digit Span Tests), executive function (increased reaction time in TMT-A and B tests) and semantic memory (poor performance in animal naming tests). A mediation model, incorporating neuroanatomic volume and PVWMH, demonstrated that PVWMH load contributes significantly to loss in executive function mediated by atrophy in a unique set of neuroanatomic structures in addition to direct impacts on cognition; while a direct effect of PVWMH is a major path for effects on working and semantic memory. Global cognition like MMSE is affected only at a higher PVWMH accumulation (>6 ml).

## Materials and Methods

### Study population and Characteristics

MR Images, clinical investigations, and cognitive status of subjects classified as cognitively normal (CN), cognitively impaired (CI), and cognitively impaired with an etiological diagnosis of Alzheimer’s disease (CI-AD) were obtained from National Alzheimer’s Coordinating Center (NACC). Subjects, aged 50 to 95 years, were enrolled from 2005 to 2021. A total of 986 subjects (first visit) were included in the study. NACC is a longitudinal multicenter study established in 1999 by the National Institute of Aging (NIA) which collects and standardizes clinical and neuropathological data from Alzheimer’s Disease Research Centers (ADRCs) across the United States (Beekly et al., 2007, 2004; Besser et al., 2018). T2-Fluid Attenuated Inversion Recovery (FLAIR) and T1-weighted (T1w) MRI from CN (N=521), CI (N=146), and CI-AD (N=319) subjects enrolled in the NACC study till September 2021 from 16 ADRCs (https://www.alz.washington.edu/), were segmented for quantification of WMH, and neuroanatomic volumetry, surface area and thickness. MRI images with slices containing the whole brain were included. Scans that lacked either T1 or T2-FLAIR modality, were excluded. Subjects were stratified across three age groups: (i) early (50-64 years), (ii) intermediate (65-79 years), and (iii) late (≥80 years) age-group as described in prior investigations (Cook et al., 2017; Geifman et al., 2013).

### Periventricular and Deep White Matter Hyperintensity segmentation and quantification

A cluster-based white matter hyperintensity (WMH) extraction pipeline based on the k-nearest neighbors (k-NN) algorithm, UBO Detector (Jiang et al., 2018), was established and optimized for WMH segmentation and quantification across the three cognitive groups: CN, CI and CI-AD. T1-weighted and T2-FLAIR MR-images were co-registered to DARTEL (Diffeomorphic Anatomical Registration Through Exponentiated Lie) space. kNN value and probability threshold were optimized before the segmentation of brain MR images to determine WMH volume in the periventricular and deep white matter regions. WMH located within a 10 mm distance from the ventricle boundary was considered as PVWMH; else, it was categorized as DWMH. The segmented total WMH lesion masks were categorized by volume as punctate (<10mm^3^), focal (10-30 mm^3^), medium (30-50 mm^3^) and confluent (>50 mm^3^) lesions. The DWMH was further quantified as frontal, temporal, parietal, and occipital- DWMH using lobar masks.

### Probability map generation to investigate WMH spatial distribution

To obtain a likelihood of WMH occurrence in periventricular and deep white matter regions, the PVWMH and DWMH masks obtained after segmentation were binarised, with ‘1’ indicating WMH presence, and ‘0’ indicating WMH absence in each voxel (with a voxel size 1x1x1 mm). A voxel-wise probability map of PVWMH and DWMH distribution was generated for the three age groups in CN, CI and CI-AD subjects. The voxel-wise probability map of PVWMH and DWMH was estimated by overlapping the respective WMH masks, summing the voxel values, and normalizing by the total number of WMH images within each age group.

### Periventricular and deep WMH kinetics with age

An exponential growth curve model was fitted and optimized for investigating the kinetics of PVWMH or DWMH progression with age across CN, CI and CI-AD groups as following:

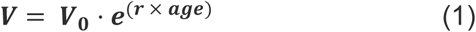

Where *V* is PVWMH or DWMH volume at the given age, *V*_0_ is initial PVWMH or DWMH volume at 50 years of age, *r* is the rate constant. The instantaneous rate of change of *V* was calculated to understand how rapidly the volume changes at a given age. This rate, represented by the derivative of PVWMH or DWMH volume with age, was determined using the formula:

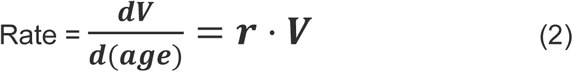

Further, to identify an inflection point in the exponential trajectory of WMH volume with age, piecewise linear least-square fitting was performed using the ’pwlf’ python library (Jekel, 2019).

### Neuroanatomical region segmentation and quantification

The cross-sectional MRI data from CN subjects (N=521) were quantified for neuroanatomical volumes and thickness by segmenting T1w MR image using standard ‘recon-all’ processing pipeline within FreeSurfer (v7.3.2) (Fischl, 2012). A total of 174 parameters were quantified including volumetry and thickness of neuroanatomic structures. Brain region volumes were normalized to the estimated Total Intracranial Volume (eTIV) using Equation (3) (Müller et al., 2019; Thota et al., 2021):

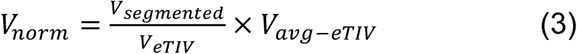

Where *V*_*norm*_ is the normalized volume; *V*_*segmented*_ is the segmented volume of brain regions, *V*_*eTIV*_ is the estimated total intracranial volume and *V*_*avg−eTIV*_ represents mean eTIV for each age-group. The average cortical thickness of neuroanatomic regions was calculated by averaging the thickness from the two hemispheres.

Further the longitudinal neuroanatomic changes were estimated in CN subjects (N=27) who had at least two consecutive visits within a maximum interval of 5 years. The change in structural volume (*ΔV*) between visits was calculated as follows:

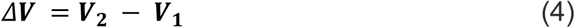

Where, *V*_1_ represents volume of brain region at the first visit, and *V*_2_ is the volume at the second visit. Longitudinal changes in structural volume between the visits were calculated for subjects within a specific range of WMH load change.

### Threshold of PVWMH and DWMH, Cognitive performance and Neuroanatomic Mediation in CN subjects

A battery of neuropsychological test scores of CN subjects was obtained from the NACC Uniform Data Set (UDS) (Morris et al., 2006), which had a corresponding MRI measurement. The global cognition was evaluated using the Mini-Mental State Examination (MMSE; Total score using D-L-R-O-W) (Folstein et al., 1975). The episodic memory function was assessed using Logical Memory IIA -Delayed (LMT, Total story units recalled) while semantic memory and verbal fluency were estimated using Animal naming test (in 60 seconds). Language abilities were examined using the Boston Naming Test (BNT). Attention and working memory were estimated using the Digit Span Test (DST) - forward and backward, while the executive function was measured using the Trail Making Test part-A and B (total number of seconds to complete the test). TMT scores reflect reaction time, with higher scores indicating poorer performance. In contrast, higher MMSE, LMT, Animal Naming, and DST scores indicate better cognitive performance.

To estimate the threshold of total WMH, PVWMH and DWMH volume leading to cognitive and neuroanatomical perturbations, CN subjects were grouped into four quartiles (Q1, Q2, Q3, Q4). Cognitive test scores were then compared across these quartiles. Only subjects with cognitive data (N=389 out of 521 CN subjects) were included in the analysis.

To examine the plausible implication of neuroanatomic volumes and thickness as a mediator for impact of WMH on cognition, a simple mediation analysis was conducted on a sample of 521 CN subjects where PVWMH or DWMH load serves as predictor (X) and cognitive test score as an outcome (Y). The mediation analysis involved investigating following paths:

i. Interaction of WMH load with neuroanatomic volume (path a),
ii. Interaction of neuroanatomic volume with cognitive performance (path b),
iii. Interaction of WMH load with cognitive performance (direct effect; path c’), and
iv. Interaction of WMH load with cognitive performance via neuroanatomic parameters (indirect effect; path ab) (**Fig. 1**).

**Figure 1.**
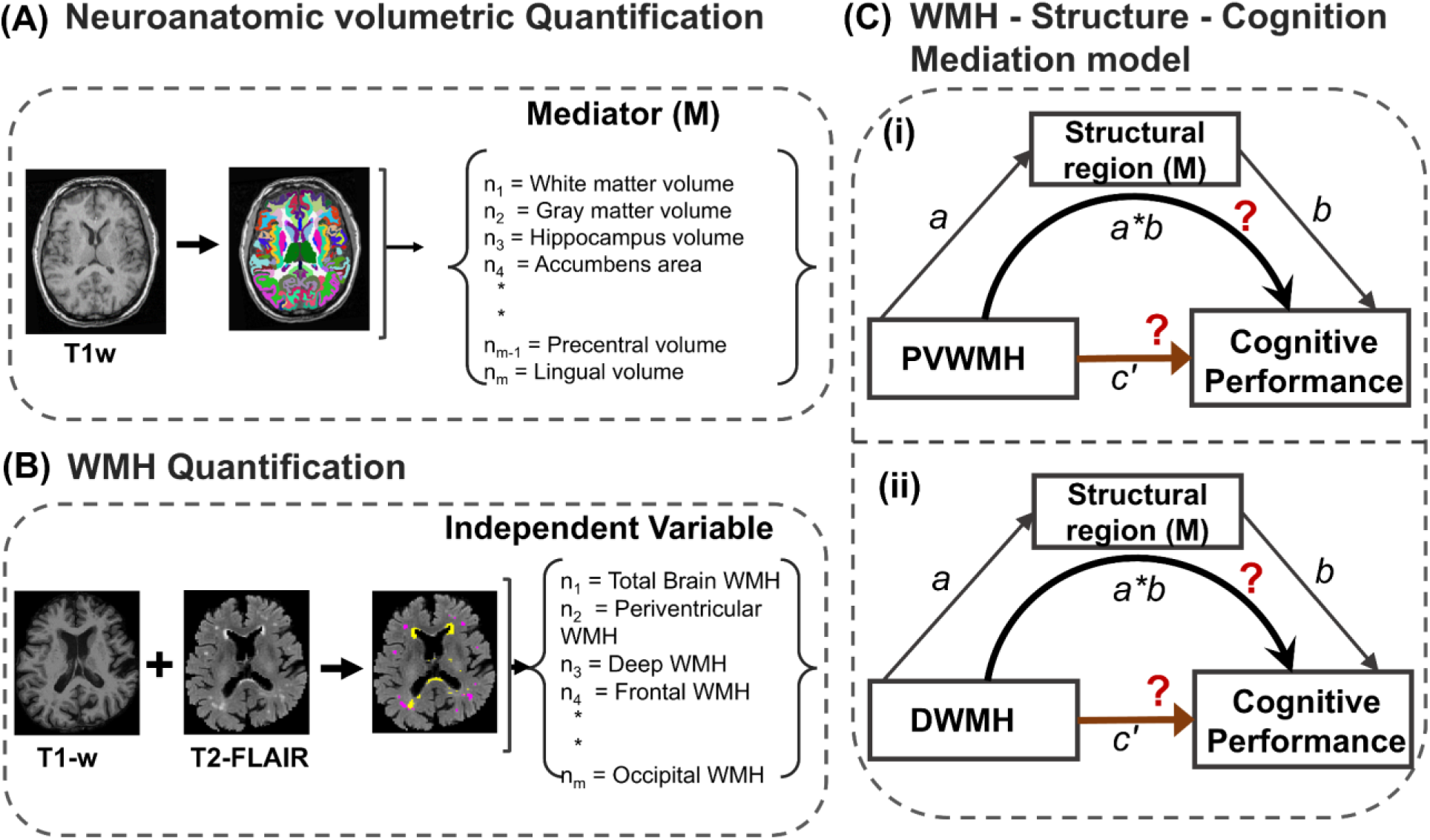
**(A)** T1w-MR images were segmented to obtain neuroanatomic features using FreeSurfer. **(B)** WMH segmentation was performed, post co-registration of T1 and T2-FLAIR modalities, to quantify regional WMH volumes. **(C)** Proposed mediation framework, where delay in reaction time (dependent variable) is mediated via atrophy of brain structural regions (mediator) due to (i) PV or (ii) Deep WMH load (independent variable).

Independent simple mediation models were run for each of 174 quantified brain structural regions as mediators. Confidence intervals were set at 95%, with 5000 bootstrap samples generated for percentile bootstrap confidence intervals to ensure robust statistical assessment. The models were adjusted for age as a covariate specified as follows:

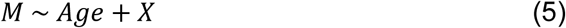

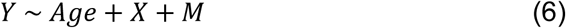

The contribution of the indirect effect of WMH on cognition towards the total effect on cognition was estimated upon dividing the coefficient of indirect effect (ab) by the coefficient of the total effect (*c′* + *ab*), multiplied by 100.

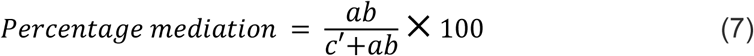

The mediation analysis was performed using Model 4 of the PROCESS tool developed by Andrew F. Hayes, implemented in the R statistical environment (Bolin, 2014).

### Statistical analysis

The groupwise differences in PVWMH and DWMH load between the three age groups across the cognitive groups (CN, CI and CI-AD) was estimated using non-parametric statistical Mann Whitney U test. The Bonferroni correction was employed to reduce the chances of obtaining type I errors while performing multiple pairwise tests.

The exponential fitting to obtain the progression rate between CN, CI and CI-AD also employed the uncertainty in estimates using a 95% CI. Instantaneous rates of change in PVWMH and DWMH volume were calculated annually from ages 50 to 95 years using exponential fitting models (*V* = *V*_0_ · *e*^(*r* × *age*)^) specific to CN, CI, and CI-AD groups. For each age group, the mean rate of increase was calculated. The rate of PVWMH relative to DWMH in the three age-groups were calculated as:

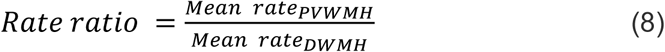

Rate of change in neuroanatomic volume with age was modeled using linear regression with the intercept adjusted to 50-years of age by subtracting 50 from the age at the respective visit, so as to account for variations at the entry of the subjects in the study.

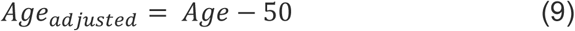

To assess the relative importance of interacting predictors (PVWMH, frontal WMH, temporal WMH, parietal WMH and occipital WMH loads) in a statistical model, a python library called Dominance-Analysis was used. The method accounts for the individual effect of each regional WMH load as well as its effect in conjunction with other regions, identifying its proportionate contribution to the model.

To understand the effect of WMH load on cognition and neuroanatomic volumetry, and to identify a threshold of PVWMH and DWMH, a one-way ANCOVA was performed using SPSS (version 29.0.2.0) , keeping age as a covariate, WMH quartiles as independent variable, and cognitive test scores as dependent variables. This analysis controlled for age-related variance to isolate the impact of WMH on cognition. Post-hoc pairwise comparisons with Bonferroni correction were applied to identify significant differences between WMH quartiles. A Wilcoxon rank-sum test was used to assess significant differences in brain neuroanatomic volumes between two longitudinal visits.

## Result

### Demographics

The cross-sectional MRI and cognitive measurements of 986 subjects from the NACC cohort was included in the study, comprising 521 cognitively normal (CN) subjects (mean age: 66 ± 8.4 years), 146 cognitively impaired (CI) subjects (mean age: 73.1 ± 9.4 years), and 319 CI-AD subjects (mean age: 76 ± 8.1 years) (**Table 1**).

**Table 1.**
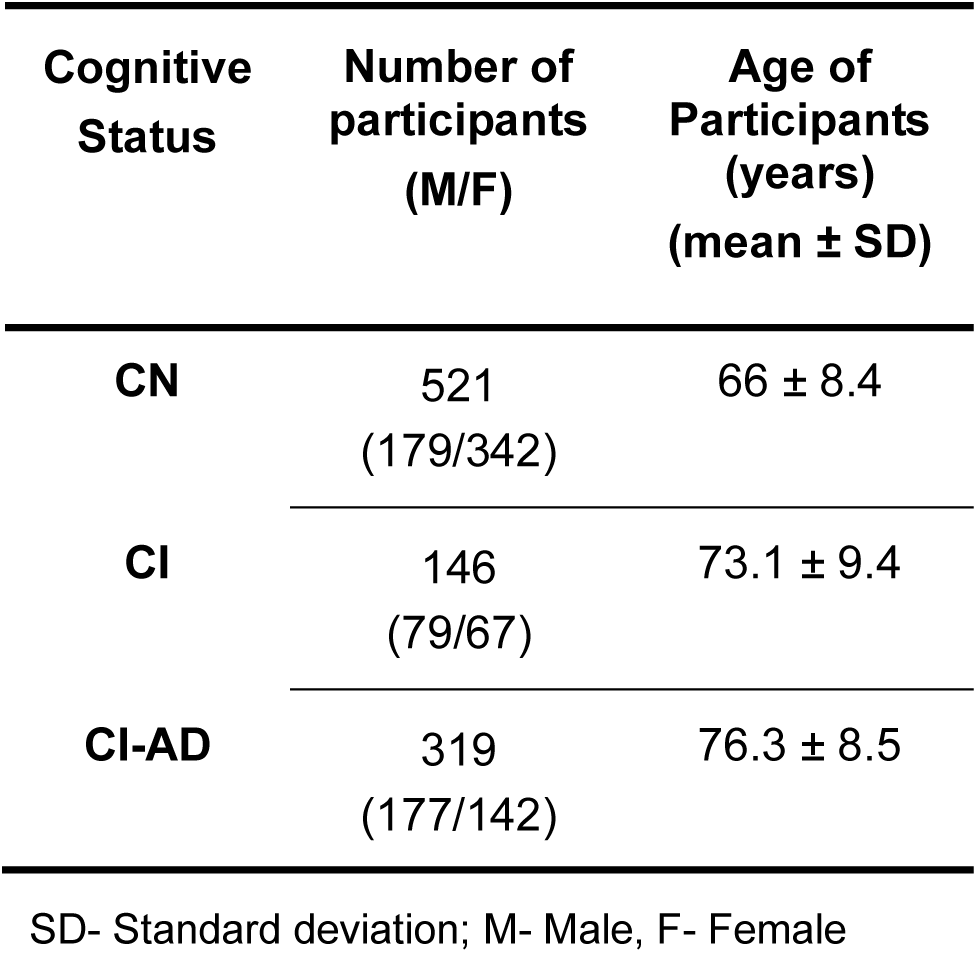
Demographics of the cohort.

| **Cognitive Status** | **Number of participants (M/F)** | **Age of Participants (years) (mean ± SD)** |
| --- | --- | --- |
| **CN** | 521<br>(179/342) | 66 ± 8.4 |
| **CI** | 146<br>(79/67) | 73.1 ± 9.4 |
| **CI-AD** | 319<br>(177/142) | 76.3 ± 8.5 |

SD- Standard deviation; M- Male, F- Female

### Spatial Probability of PVWMH and DWMH with Age in CN, CI and CI-AD

WMH segmentation revealed that ∼50% of the CN subjects in the early age group presented with PVWMH and DWMH load more than 1 ml, while ∼85% of subjects in the intermediate age and >90% of the subjects in the late age groups exhibited significant PVWMH and DWMH load (>1 ml). The prevalence of PVWMH and DWMH load increased significantly in CI and CI-AD subjects at all the three age groups (**Fig. 2A, B**). The voxel-wise probability map derived from PVWMH quantification in CN subjects depicts a higher likelihood of PVWMH deposition in the ventricular horns (probability > 0.5) (**Fig 2C**), with a significantly greater probability of WMH presence in the frontal horn compared to the occipital horn (p < 0.01). Noticeably, the probability of WMH in the frontal and occipital horns of the CI and CI-AD groups was ∼2-3 times higher compared to the CN group at the early and intermediate age groups. The probability of DWMH for all the three cognitive groups was diffused and lower compared to PVWMH even at the early age groups indicative of PVWMH as an earlier incidence than DWMH (**Fig 2D**).

**Figure 2.**
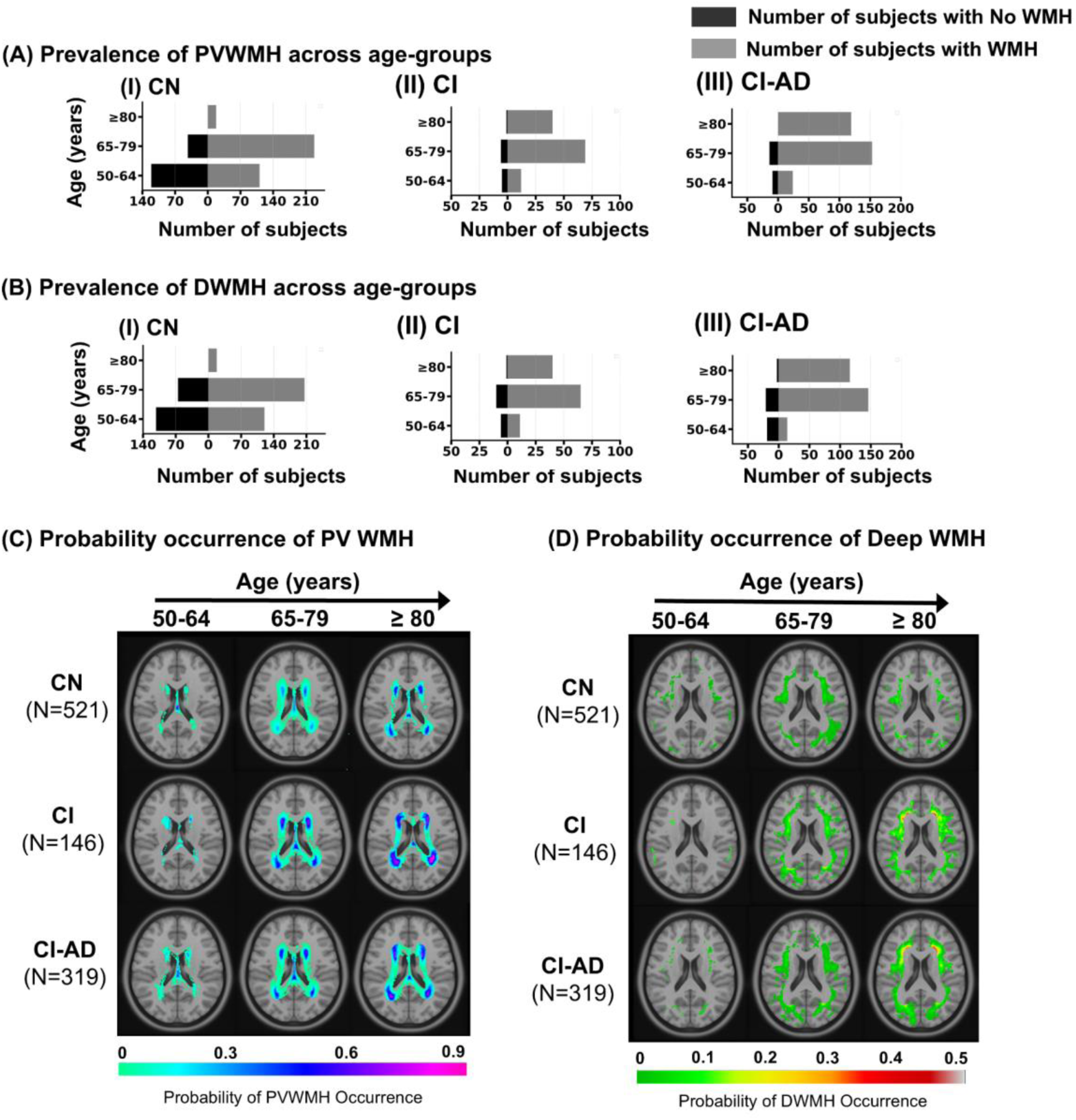
Prevalence and probabilistic distribution of white matter hyperintensities (WMH) in various age groups. Prevalence of **(A)** PVWMH and **(B)** DWMH in cognitively normal (CN), cognitively impaired (CI), and cognitively impaired with Alzheimer’s disease (CI-AD) subjects, categorized by age groups: early (50-64 years), intermediate (65-79 years), and late (>80 years). Subjects with a minimum WMH load of 1 ml are classified as having ’No WMH’. Voxel-wise probabilistic maps of WMH occurrence for **(D)** periventricular and **(E)** deep WMH, overlaid on a T1-weighted MNI template. Colormap represents probability of WMH occurrence. Higher probability values indicate regions with increased WMH deposition with aging. The study population consisted of 521 CN (N=232 early, N=271 intermediate, N=18 late), 146 CI (N=30 early, N=75 intermediate, N=41 late), and 319 CI-AD (N=33 early, N=167 intermediate, N=119 late) subjects.

### Load of PVWMH and DWMH with Age in CN, CI and CI-AD

A global statistical analysis of PVWMH load for the three age groups between the cognitive groups revealed significant differences (H = 423.99, dfbetween= 8, dfwithin= 977, p<0.001). Similarly, for DWMH load, significant differences were observed (H = 261.29, dfbetween= 8, dfwithin= 977, p<0.001). Noticeably, at the early age, CI and CI-AD subjects exhibited a significantly higher PVWMH volume compared to CN subjects (CN: 1.41 ± 1.9 ml, CI: 3.96 ± 4.7 ml, CI-AD: 2.47 ± 2 ml; p(CN vs CI) <0.001, p(CN vs CI-AD) =0.0001, and p(CI vs CI-AD) =0.28), with no significant difference between CI and CI-AD groups (**Fig. 3B**, **Table 2**). With progression in age, the PVWMH volume increased ∼3-folds for all the three cognitive groups compared to the PVWMH load at the early age groups. Similar to the pattern observed at the early age groups, PVWMH in CI and CI-AD groups compared to CN was elevated at intermediate (CN: 6.59 ± 7.8 ml, CI: 9.9 ± 8.3 ml, CI-AD: 9.39 ± 8.5 ml; p(CN vs CI) =0.0002, p(CN vs CI-AD) <0.001, and p(CI vs CI-AD) =0.59) and late age groups (CN: 7.51 ± 6.1 ml, CI: 16.43 ± 10.7 ml, CI-AD: 15.9 ± 10.2 ml; p(CN vs CI) =0.0008, p(CN vs CI-AD) =0.0002, and p(CI vs CI-AD) =0.77) while PVWMH load between CI and CI-AD was not distinct.

**Figure 3.**
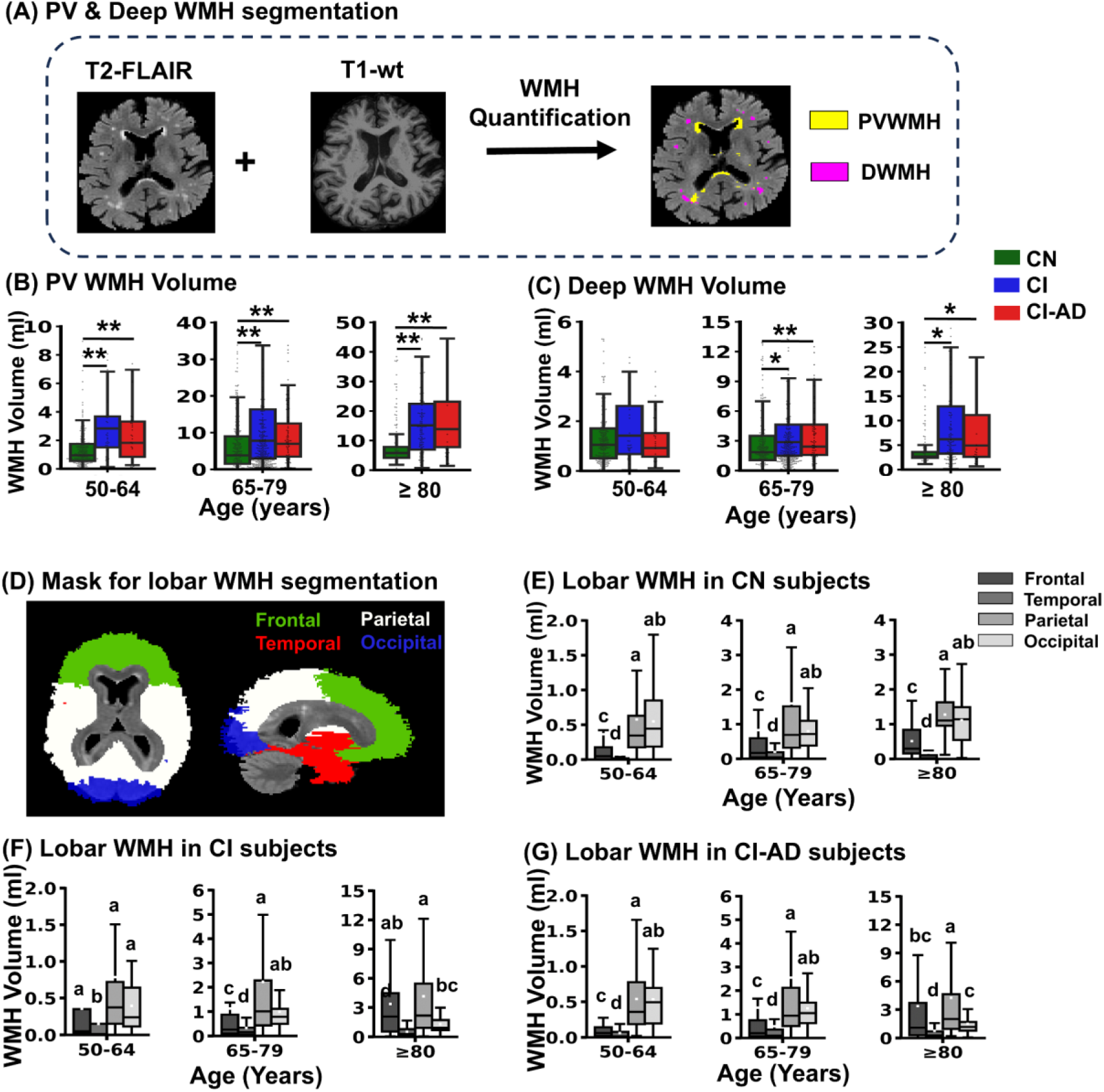
Age-related dynamics of PVWMH, DWMH and lobar WMH accumulation (A) WMH segmentation process using T1 and T2-FLAIR modalities to segment PV and DWMH. Volume distribution of WMH in the **(B)** periventricular and **(C)** deep white matter regions with aging across cognitively normal (CN), cognitively impaired (CI), and cognitively impaired with Alzheimer’s disease etiology (CI-AD) groups. **(D)** Mask used for lobar segmentation (frontal, temporal, parietal, and occipital regions). Lobar WMH volume distribution with age in **(E)** CN, **(F)** CI, and **(G)** CI-AD. The statistical differences were assessed using Mann-Whitney U tests followed by Bonferroni correction. The upper margin of the box represents the third quartile (Q3), the lower margin represents the first quartile (Q1), and the height of the box represents the interquartile range (IQR), with the median indicated by solid line. *p < 0.016, **p < 0.001. Different letters indicate statistically significant differences between lobar WMH loads. The study population consisted of 521 CN (N=232 early, N=271 intermediate, N=18 late), 146 CI (N=30 early, N=75 intermediate, N=41 late), and 319 CI-AD (N=33 early, N=167 intermediate, N=119 late) subjects.

**Table 2.** Age-groupwise WMH volume distribution across different cognitive groups.

| **Cognitive Status** | **Age group (years)** | **Total WMH volume (mean±SD)** | **PVWMH volume (mean±SD)** | **DWMH volume (mean±SD)** |
| --- | --- | --- | --- | --- |
| **CN** | **50-64** | 3.06 ± 3.6 | 1.41 ± 1.9 | 1.56 ± 2.2 |
|  | **65-79** | 10.32 ± 12.5 | 6.58 ± 7.8 | 3.42 ± 5.3 |
|  | **≥80** | 10.81 ± 7.1 | 7.51 ± 5.9 | 3.08 ± 1.4 |
| **CI** | **50-64** | 4.83 ± 4.8 | 3.96 ± 4.6** | 1.68 ± 1.7 |
|  | **65-79** | 14.94 ± 12.9 | 9.9 ± 8.2** | 4.66 ± 5.6* |
|  | **≥80** | 26.36 ± 17.9 | 16.43 ± 10.5** | 9.32 ± 8.3** |
| **CI-AD** | **50-64** | 4.1 ± 3.24 | 2.46 ± 2** | 1.44 ± 1.4 |
|  | **65-79** | 15.3 ± 22.6 | 9.39 ± 8.5** | 5.56 ± 16.7* |
|  | **≥80** | 26.34 ± 2 | 15.86 ± 10.2** | 9.87 ± 15.3* |

\*Significantly different from CN (\*p<0.016, \*\*p<0.001)
SD- Standard deviation

However, DWMH load between CN, CI and CI-AD was not significantly distinct at the early age group (CN: 1.56 ± 2.2 ml, CI: 1.68 ± 1.7 ml, CI-AD: 1.44 ± 1.5 ml; p(CN vs CI) =0.053, p(CN vs CI-AD) =0.86, and p(CI vs CI-AD) =0.1) unlike the PVWMH load. But, at intermediate and late age groups, CI and CI-AD presented with significantly higher DWMH load compared to CN while CI and CI-AD had similar load (**Fig. 3C**, **Table 2**). Further, quantification of DWMH across frontal, parietal, temporal, and occipital regions revealed lobar variations of WMH across the cognitive groups. In the CN subjects, WMH load was highest in Parietal lobe ∼ Occipital lobe > Frontal lobe > temporal lobe at all the age groups (**Fig. 3E**). Similar to the CN subjects, the CI and CI-AD subjects exhibited the similar pattern of WMH load during early and intermediate age groups. However, in the late age groups, CI and CI-AD subjects exhibited an elevated WMH load in the frontal lobe, followed by the parietal lobe ∼ occipital lobe > temporal lobe. (**Fig. 3F, G**).

### Kinetics of PVWMH and DWMH with Age in CN, CI and CI-AD

Across all the three cognitive groups; increase in volume of PVWMH and DWMH with age followed an exponential growth pattern. The rate of increase of PVWMH was 1.9 ± 0.2- times faster than the increase in DWMH (rPVWMH: 0.57 ± 0.5; rDWMH: 0.18 ± 0.1) at the early age group in the CN subjects. At the intermediate age group, the rate of increase in PVWMH relative to DWMH was 2.58 ± 0.2 times-faster and it increased to 3.57 ± 0.3 times in the late age group (**Fig 4A, D, Supp. Table 1**). For CI subjects in the early age group, the rate of PVWMH increase was 1.86 ± 0.02 times faster than DWMH (rPVWMH: 0.48 ± 0.03; rDWMH: 0.27 ± 0.02). This relative rate of increase for PVWMH compared to DWMH remained consistent across the intermediate and late age groups, with ratios of 1.79 ± 0.02 and 1.73 ± 0.02, respectively (**Fig 4B, D**). While in the CI- AD subjects, PVWMH increased approximately 1.66 ± 0.01 times faster (rPVWMH: 0.47 ± 0.3; rDWMH: 0.29 ± 0.2) than DWMH across all the age groups (**Fig 4C, D**).

**Figure 4.**
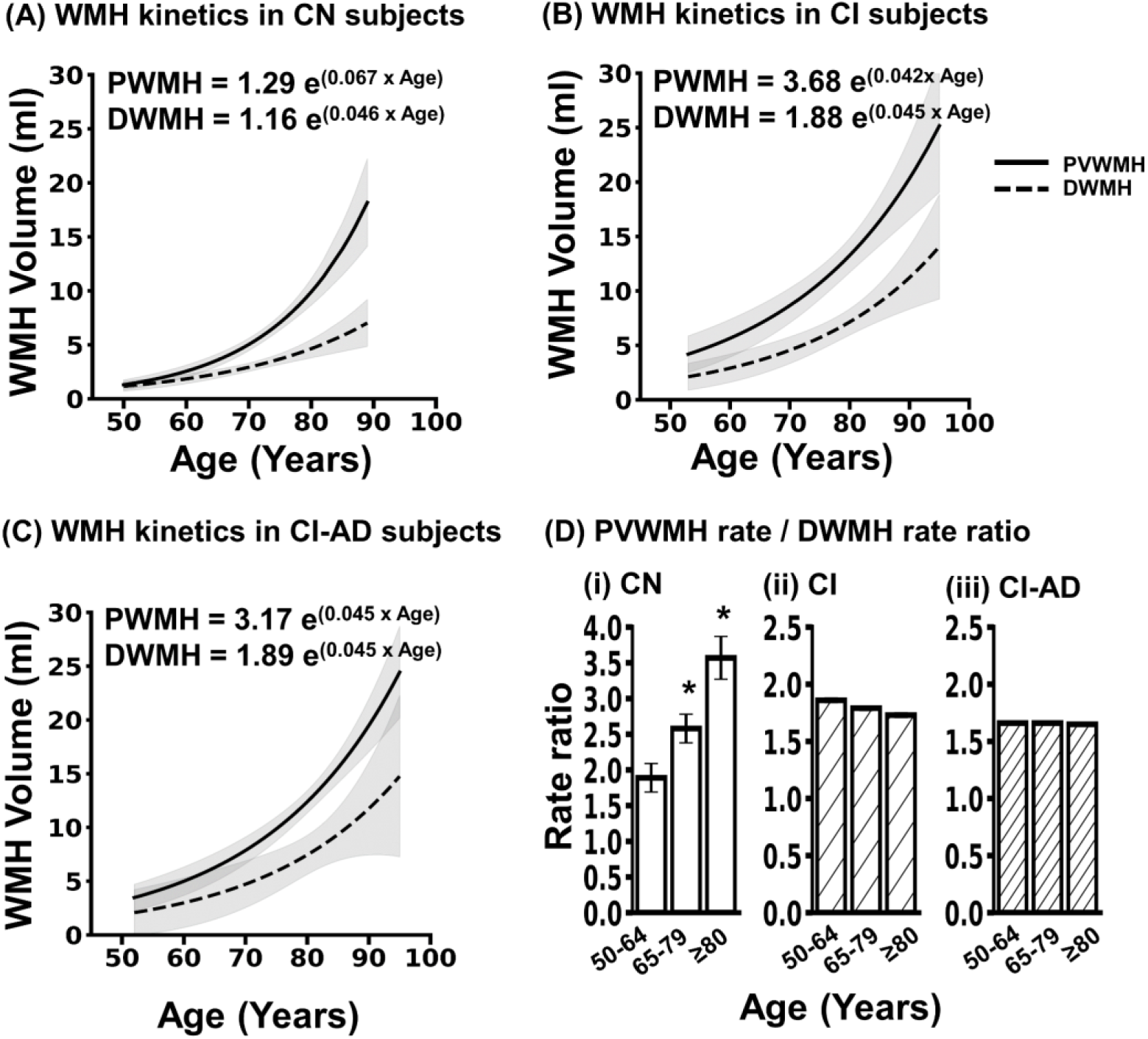
Age-related dynamics of PV and Deep WMH accumulation. The burden of PVWMH (solid line) and DWMH (dashed line) increases exponentially with age across **(A)** CN, **(B)** CI, and **(C)** CI-AD subjects. The equation represents the rate of change of WMH load where, r is the rate constant and V0 is the initial WMH volume at 50 years of age. Volume of WMH with age is fitted with a mono-exponential model in Python using *scipy* package, *curve_fit* and *kmfit.* The shaded gray area represents a 95% confidence interval. **(D)** Bar plots represent the mean rate ratio of PVWMH and DWMH growth for early, intermediated and late age groups in CN, CI and CI-AD groups. *p-value<0.001

The progression of WMH modeled using piecewise linear regression revealed that the total WMH volume remained stable in the age range of 50 to 60 years with a slope of β = 0.09 (p = 0.07) in CN subjects. An inflection point was identified at 61 years of age beyond which a significant escalation in total WMH was observed (β = 0.64, p < 0.001). Similarly, PVWMH volume remained low and constant until approximately 61.2 years (β = 0.02, p = 0.46), after which it increased significantly (β = 0.48, p < 0.001), indicating a knee-point in PVWMH progression kinetics. However, DMWH kinetics did not present a distinct inflection point with age (**Supp.** Fig. 1).

### WMH lesion distribution with age

Further segmentation of PVWMH and DWMH lesions based on lesion volume categorized into: punctate (<10 mm³), focal (10-30 mm³), medium (30-50 mm³) and confluent (>50 mm³) (**Fig. 5A**). In CN subjects, at early and intermediate ages, the order of lesion frequency was punctate > focal ≈ confluent > medium, with medium-sized lesions showing minimal increase with age (**Fig. 5B**). In deep white matter, the hierarchy was focal > punctate ≈ confluent > medium at early age; whereas at intermediate and late ages, it was punctate ≈ focal > confluent > medium (**Fig. 5C**). Notably, the number of all lesion categories increased significantly with age (p < 0.001). The age-groupwise distribution of lesion types in periventricular and deep white matter for CN, CI and CI- AD subjects is detailed in **Table 3A** and **3B**, respectively.

**Figure 5.**
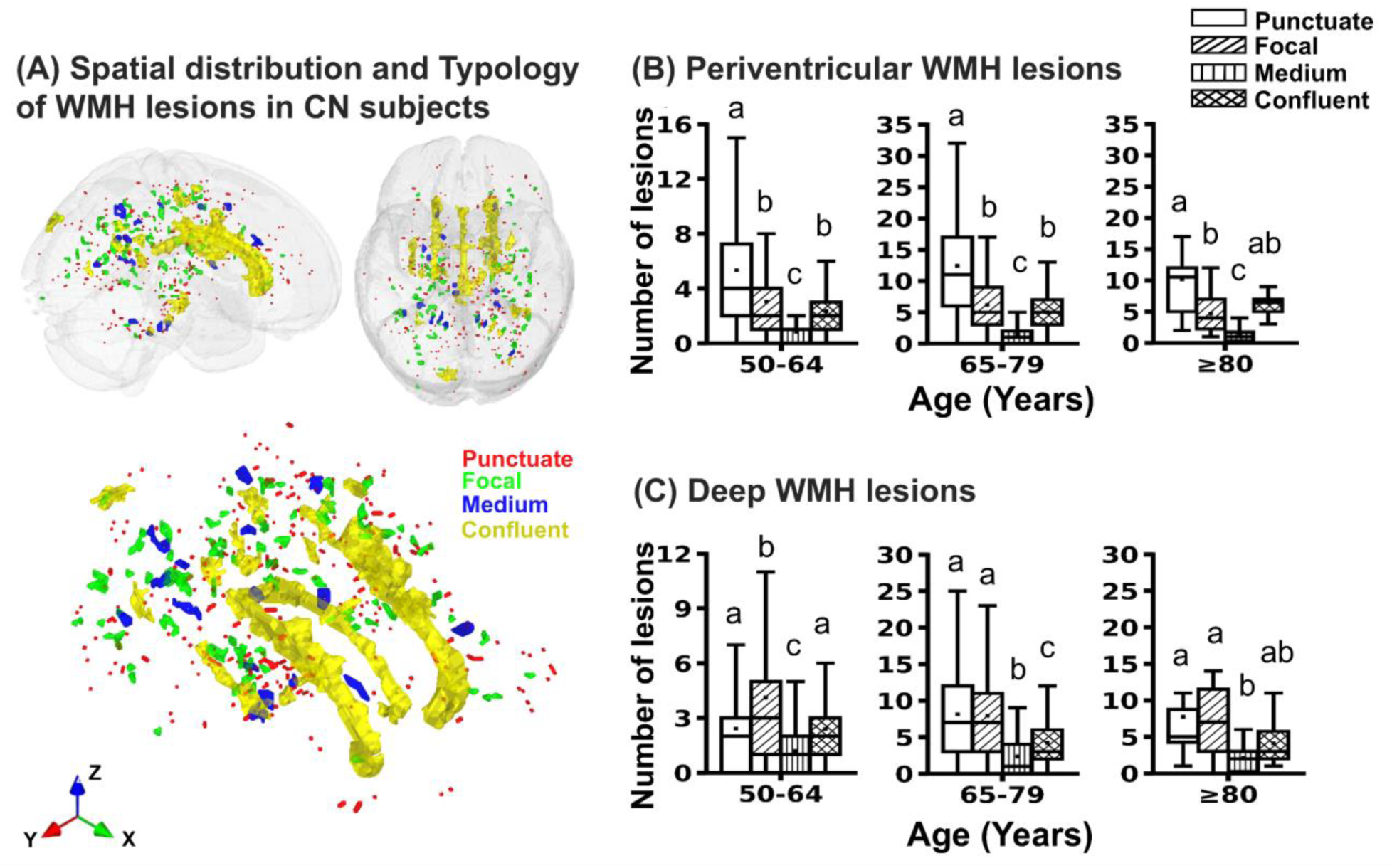
Age-Related Distribution and Quantification of WMH Lesions by Type (A) Representative WMH lesion mask annotation based on lesion volume: punctate (< 10 mm3 i.e. 3 voxels on DARTEL space; depicted in red), focal (10 - 30 mm3, i.e. 9 voxels; depicted in green), medium (30 - 50 mm3 i.e. 15 voxels; depicted in blue) and confluent (> 50 mm3; depicted in yellow). Quantification of number of lesions with age in **(B)** Periventricular region, **(C)** Deep white matter region in cognitively normal subjects (N=521). The statistical differences were tested using Mann-Whitney U tests followed by Bonferroni correction. The upper margin of the box represents the third quartile (Q3), the lower margin represents the first quartile (Q1), and the height of the box represents the interquartile range (IQR), with the median indicated by solid line, and mean as white square. Different letters indicate statistically significant differences between the lesion types (p<0.012).

**Table 3A.** Age-groupwise distribution of punctate, focal, medium and confluent PVWMH lesions across different cognitive groups.

| **Cognitive Status** | **Age group (years)** | **Punctate (mean ± SD)** | **Focal (mean ± SD)** | **Medium (mean ± SD)** | **Confluent (mean ± SD)** |
| --- | --- | --- | --- | --- | --- |
| **CN** | **50-64** | 5.3 ± 4.8 | 3 ± 2.7 | 0.8 ± 1.1 | 2.3 ± 1.8 |
| | **65-79** | 12.4 ± 8.1 | 6.2 ± 4.3 | 1.5 ± 1.5 | 5 ± 2.8 |
| | **$\geq 80$** | 10.1 ± 6.2 | 4.7 ± 2.8 | 1.3 ± 1.2 | 6.4 ± 1.8 |
| **CI** | **50-64** | 10.1 ± 10 | 4.8 ± 4.47 | 1.2 ± 1.8 | 3.7 ± 4 |
| | **65-79** | 12.8 ± 9.53 | 5.4 ± 3.6 | 1.3 ± 1.3 | 4.8 ± 2.5 |
| | **$\geq 80$** | 14.8 ± 7.8 | 5.8 ± 3.66 | 1.4 ± 1.37 | 5.4 ± 2.4 |
| **CI-AD** | **50-64** | 10.4 ± 7.1 | 5.6 ± 4.3 | 1.8 ± 1.6 | 4.1 ± 2.9 |
| | **65-79** | 11 ± 6.5 | 5.1 ± 3.1 | 1.3 ± 1.4 | 5.4 ± 2.5 |
| | **$\geq 80$** | 13.6 ± 8.5 | 5.4 ± 3.9 | 1.5 ± 1.6 | 6.1 ± 2.4 |

**Table 3B.**
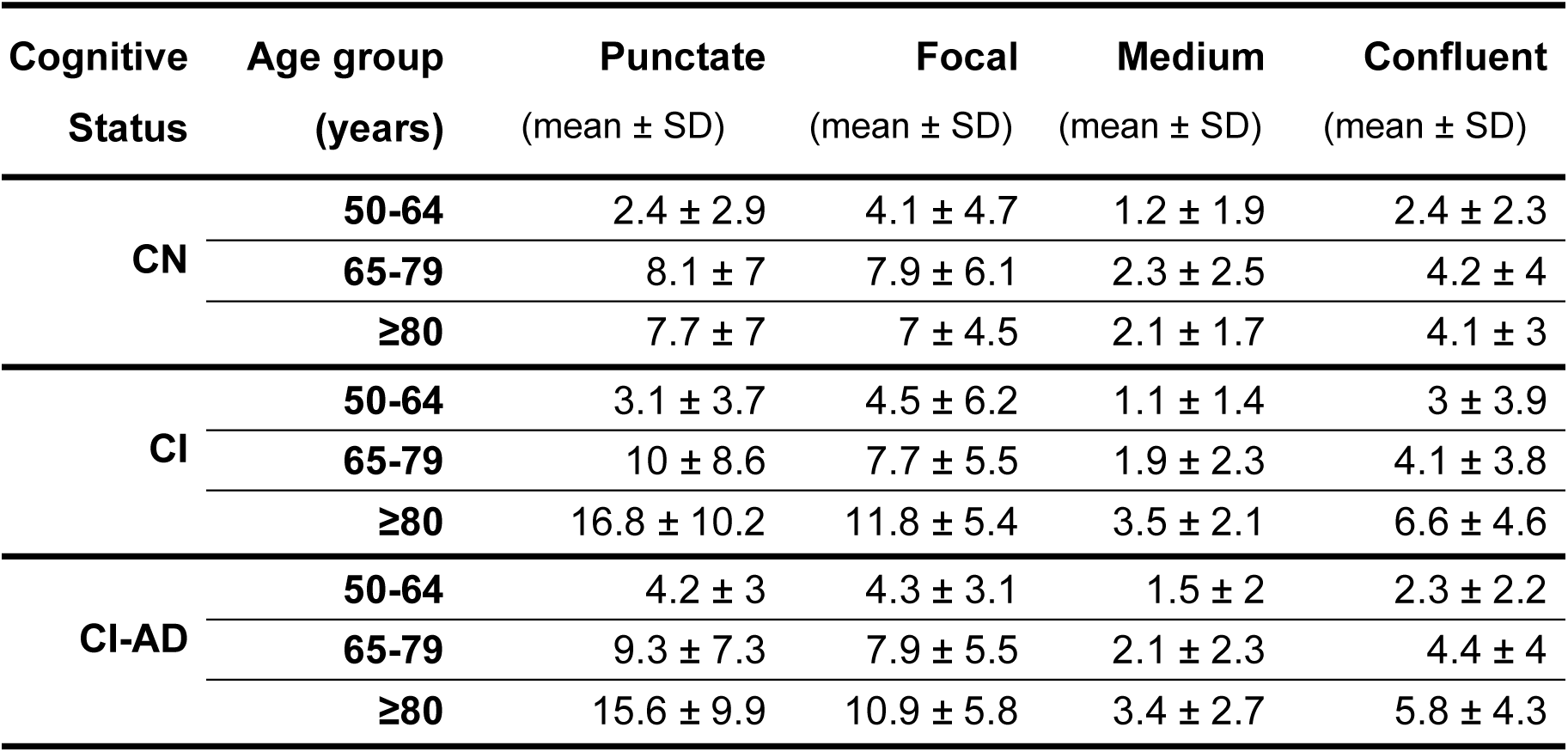
Age-groupwise distribution of punctate, focal, medium and confluent DWMH lesions across different cognitive groups.

### Impact of PVWMH and DWMH on Cognitive Domains with Age

Multivariate linear regression model examining the association between cognitive performances, age, PVWMH and DWMH load revealed a positive association between increase in PVWMH load and reaction time in TMT-A (β=0.40, p= 0.01) and TMT-B (β=1.86, p=0.001). Negative associations were observed between PVWMH and cognitive scores of DST-B (β=-0.05, p= 0.04) animal naming (β=-0.20, p= 0.003) (**Table 4**). No significant associations were found between DWMH load and cognitive scores.

**Table 4.** Multivariate linear regression model coefficients and standard error for cognitive test scores.

| Cognitive tests | $\beta 0$ | | $\beta 1$ | | $\beta 2$ | | $\beta 3$ | |
| --- | --- | --- | --- | --- | --- | --- | --- | --- |
|  | Value | SE | Value | SE | Value | SE | Value | SE |
| **MMSE** | 29.7** | 0.15 | -0.04** | 0.01 | -0.03 | 0.02 | 0 | 0.02 |
| **BNT** | 29.16* | 0.29 | -0.08** | 0.02 | -0.05 | 0.03 | 0.05 | 0.04 |
| **LMT** | 13.91** | 0.44 | -0.07** | 0.03 | -0.09 | 0.05 | 0 | 0.07 |
| **Animal naming** | 25.6** | 0.6 | -0.18** | 0.04 | -0.20** | 0.06 | 0.09 | 0.09 |
| **DST-F** | 9.59** | 0.23 | -0.06** | 0.01 | -0.03 | 0.02 | -0.01 | 0.03 |
| **DST-B** | 7.9** | 0.25 | -0.05** | 0.02 | -0.05* | 0.03 | -0.04 | 0.04 |
| **TMT-A** | 18.76* | 1.38 | 0.64** | 0.08 | 0.40** | 0.15 | -0.43 | 0.21 |
| **TMT-B** | 36.52** | 5.37 | 2.43** | 0.32 | 1.86** | 0.58 | -0.8 | 0.81 |

\*p<0.05, \*\*p<0.01
$\beta 0$ - Intercept, $\beta 1$ – coefficient for Age, $\beta 2$ – coefficient for PVWMH, $\beta 3$ – coefficient for DWMH, SE – Standard error in the estimated values of the coefficients

Stratification of the CN subjects in four quartiles based on WMH load provided a range of **Total WMH** (Q1: ≤ 2.16 ml, Q2: 2.17 - 4.14 ml, Q3: 4.15 - 8.79 ml, and Q4: >8.79 ml), **PVWMH** (Q1: ≤ 0.93 ml, Q2: 0.94 - 2.32 ml, Q3: 2.33 - 6.12 ml, and Q4: >6.12ml) and **DWMH** (Q1: ≤ 0.92 ml, Q2: 0.93 - 1.54 ml, Q3: 1.55 - 2.75 ml, Q4: >2.75 ml). Q1, Q2, Q3, and Q4 quartiles comprised 98, 97, 97, and 97 CN subjects, respectively (**Fig. 6A**).

**Figure 6.**
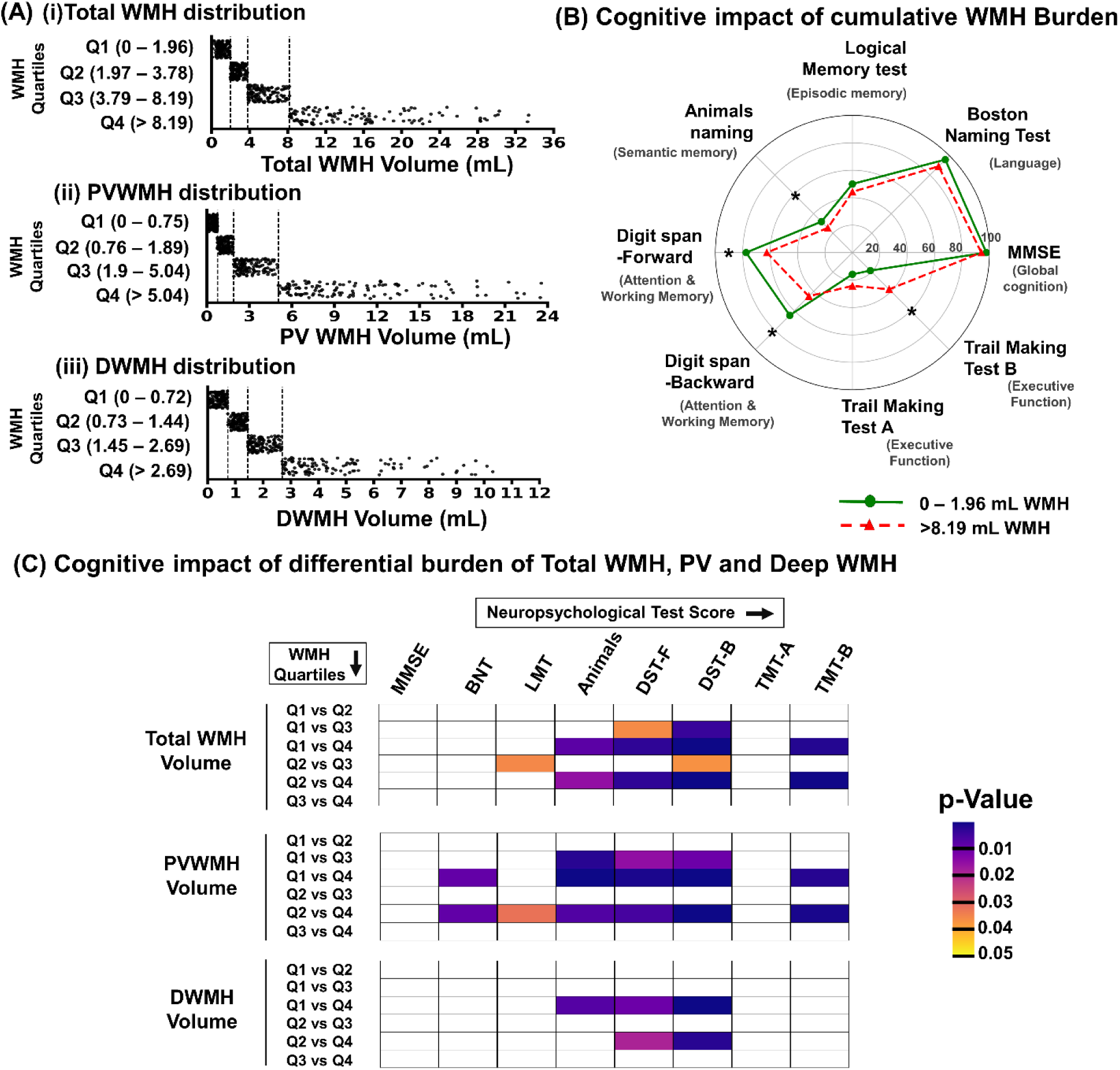
Cognitive deficits in different cognitive domains with increase in WMH (A) CN Subjects (N=389) divided into quartiles based on (i) total WMH volumes (≤ 2.16 ml, 2.17 - 4.14 ml, 4.15 - 8.79 ml, and >8.79 ml) (ii) PVWMH (≤ 0.93 ml, 0.94 - 2.32 ml, 2.33 - 6.12 ml, and >6.12ml); and (iii) DWMH volume (≤ 0.92 ml, 0.93 - 1.54 ml, 1.55 - 2.75 ml, >2.75 ml). All WMH quartiles (Q1, Q2, Q3, Q4) have sample sizes of 98, 97, 97, 97 subjects, respectively. **(B)** Radar chart represents the group means for subjects with total WMH volume <2.16 ml (green circle); and >8.79 ml (red triangle) of the neuropsychological test scores converted to percentages against the maximum score for each test. TMT scores correspond to the time (in seconds) taken to complete the test, wherein a higher score indicates poorer cognitive performance. **(C)** Heatmap represents significant differences between two given WMH quartile groups for various age-adjusted neuropsychological tests. Colormap represents the significance (p-value < 0.05). The boxes represent differences in cognitive performance across WMH quartiles for the following tests: MMSE (Mini Mental State Examination), BNT (Boston Naming Test), Logical Memory test IIA - Delayed (LMT), Animal naming (in 60 seconds), DST-F, DST-B (Digit Span Test) and TMT-A, TMT-B (Trail Making test A and B). Test scores were adjusted for age using ANCOVA, keeping age as covariate, cognitive scores as dependent variable and WMH quartiles as fixed factor. Post- hoc pairwise comparisons with Bonferroni correction were applied to identify significant differences between WMH quartiles.

The ANCOVA analysis for evaluation of effect of WMH load on cognition in the CN group, using age as a covariate, total WMH quartiles as independent variables, and cognitive test scores as dependent variables, demonstrated a significant effect of total WMH load on cognitive performances (scores) in TMT-B (F(3, 384) = 5.17, p = 0.002), DST-F (F(3, 384) = 6.2, p < 0.001), DST-B (F(3, 384) = 10.25, p < 0.001), LMT (F(3, 384) = 3.9, p = 0.009), and Animal naming (F(3, 384) = 4.45, p = 0.004) tests. Similarly, a global ANCOVA analysis using PVWMH quartiles and cognitive scores demonstrated significant impacts of PVWMH on cognitive performances in TMT-A, TMT-B, DST-F, DST-B, LMT, BNT, MMSE, and Animal Naming tests, while DWMH quartiles showed significant changes in TMT-B, DST-F, DST-B, and Animal Naming tests. Age-adjusted cognitive test scores across the quartiles of PVWMH and DWMH are presented in **Table 5A** and **5B**, respectively.

**Table 5A.** Age-adjusted mean cognitive scores across PVWMH quartiles groups (N=389)

| Cognitive domains | Neuro-psychological tests | Q1<br>($\leq 0.93$ mL) | Q2<br>(0.94 - 2.32 mL) | Q3<br>(2.33 - 6.12 mL) | Q4<br>($>6.12$ mL) |
| --- | --- | --- | --- | --- | --- |
| Global Cognition | MMSE | 29.11 ± 0.15 | 29.06 ± 0.14 | 28.82 ± 0.13 | 28.44 ± 0.15 * |
| Language | Boston Naming Test | 28.41 ± 0.28 | 27.98 ± 0.26 | 27.54 ± 0.26 | 26.97 ± 0.29 * |
| Episodic memory | Logical Memory Test | 12.65 ± 0.44 | 13.39 ± 0.41 | 11.55 ± 0.4 | 11.43 ± 0.45 |
| Semantic memory | Animals Naming Test | 23.98 ± 0.59 | 22.4 ± 0.55 | 21.09 ± 0.54 * | 20.25 ± 0.61 * |
| Executive Function | DST-Forward | 9.23 ± 0.22 | 8.67 ± 0.2 | 8.08 ± 0.2 * | 7.67 ± 0.23 * |
| | DST-Backward | 7.57 ± 0.25 | 7.13 ± 0.23 | 6.27 ± 0.23 * | 5.62 ± 0.26 * |
| | TMT-A | 27.56 ± 1.36 | 27.7 ± 1.26 | 33.42 ± 1.25 * | 31.9 ± 1.4 |
| | TMT-B | 70.68 ± 5.32 | 71.21 ± 4.95 | 91.88 ± 4.89 * | 101.3 ± 5.47 * |

N - Number of subjects; MMSE - Mini Mental State Examination; TMT - Trail Making Test; DST- Digit Span Test
\*Significantly different from Q1 (p-value<0.05)

**Table 5B.** Age-adjusted mean cognitive scores across DWMH quartiles groups (N=389)

| **Cognitive domains** | **Neuro-psychological tests** | **Q1**<br>**($\leq 0.92$ mL)** | **Q2**<br>**(0.93 - 1.54 mL)** | **Q3**<br>**(1.55 - 2.75 mL)** | **Q4**<br>**($>2.75$ mL)** |
| --- | --- | --- | --- | --- | --- |
| **Global Cognition** | **MMSE** | 28.79 ± 0.14 | 28.97 ± 0.13 | 29.09 ± 0.13 | 28.58 ± 0.14 |
| **Language** | **Boston Naming Test** | 27.91 ± 0.27 | 27.69 ± 0.26 | 27.99 ± 0.26 | 27.32 ± 0.27 |
| **Episodic memory** | **Logical Memory Test** | 11.76 ± 0.42 | 12.5 ± 0.4 | 12.83 ± 0.4 | 11.94 ± 0.43 |
| **Semantic memory** | **Animals Naming Test** | 22.99 ± 0.57 | 22.01 ± 0.54 | 21.13 ± 0.54 | 20.59 ± 0.57 * |
| **Executive Function** | **DST-Forward** | 8.72 ± 0.21 | 8.72 ± 0.2 | 8.42 ± 0.2 | 7.8 ± 0.22 * |
| | **DST-Backward** | 7.03 ± 0.24 | 7.11 ± 0.23 | 6.58 ± 0.23 | 5.86 ± 0.24 * |
| | **TMT-A** | 29.48 ± 1.31 | 29.94 ± 1.25 | 28.71 ± 1.25 | 32.42 ± 1.32 |
| | **TMT-B** | 75.73 ± 5.15 | 86.52 ± 4.91 | 79.04 ± 4.91 | 93.72 ± 5.19 |

N - Number of subjects; MMSE - Mini Mental State Examination; TMT - Trail Making Test; DST- Digit Span Test
\*Significantly different from Q1 (p-value<0.05)

A radar plot comparing the percentage cognitive scores between subjects with total WMH load in Q1 (≤ 2.16 ml) and Q4 (>8.79 ml) revealed significantly impaired performance in TMT-B (-38.3%, p = 0.009), DST-F (-14.6%, p = 0.002), DST-B (-26.2%, p < 0.001), and Animal Naming (-14.6%, p < 0.001) for the subjects with WMH load in Quartile 4 (**Fig. 6B**). However, a deeper examination revealed that cognitive scores are affected only in the WMH range of a particular quartile and beyond. A total WMH load threshold >4.2 ml (Quartile Q3) was associated with significant decline in DST-F (-11.8%, p = 0.006), DST-B (-18.1%, p = 0.001), and Animal Naming (-9.9%, p = 0.04) (**Fig. 6C**).

Comparison of cognitive performances across PVWMH quartiles revealed significant impairments for WMH load beyond a threshold of PVWMH load >2.3 ml i.e. Q3: 2.3 - 6.1 ml. The subjects in Q3 quartile showed significant impairments in DST-forward (-12.4%, p = 0.002), DST-backward (-17.2%, p = 0.002), TMT-A (-21.3%, p = 0.015), TMT-B (-29.9%, p = 0.03) and animal naming test (-12%, p = 0.004) compared to subjects in Q1. The PVWMH load >6.1 mL i.e. Q4 resulted in significant deficits in BNT (-5%, p = 0.008), LMT (-9.6%, p = 0.01) and MMSE (-2.3%, p = 0.02) tests which were not observed for subjects in Q3. However, no significant effects were observed in cognitive performances in TMT-A, TMT-B, LMT, BNT, and MMSE across any of the DWMH quartiles (**Fig. 6C**). Noticeably, beyond a threshold of DWMH load >2.75 ml (Q4: >2.75 mL), a significant impairment was observed in DST-forward (-10.5%, p = 0.028), DST-backward (-16.6%, p = 0.008) and animal naming tests (-10.4%, p = 0.03).

### PVWMH Impacts the Cognitive Domains via Unique Set of Neuroanatomic Changes

The simple mediation model analysis evaluating the direct and indirect effect of PVWMH on cognition via 174 Neuroanatomic structures showed that PVWMH load is positively associated with TMT-B completion time (β = 1.86, p = 0.001) and it is mediated via atrophy in a unique set of 6-neuroanatomical structures. Mediation analysis at the three age groups showed that a significant structural mediation is observed only at the intermediate age group (65-79 years; N=271).

PVWMH-induced cognitive deficit in TMT-B was found to be significantly mediated by brain regions associated with executive functions namely the rostral middle frontal gyrus (RMFG) (Effect = 0.45, SE = 0.17, p = 0.008, 95% CI = 0.18, 0.83), and lingual gyrus (Effect = 0.43, SE = 0.15, p = 0.004, 95% CI = 0.17, 0.76) with mediation proportions of 31% and 30% respectively. Additionally, total gray matter volume demonstrated a significant indirect effect (43.1%) on TMT-B performance (Effect = 0.61, SE = 0.22, p = 0.007, 95% CI = 0.26, 1.17) (**Fig. 7**, **Table 6**).

**Figure 7.**
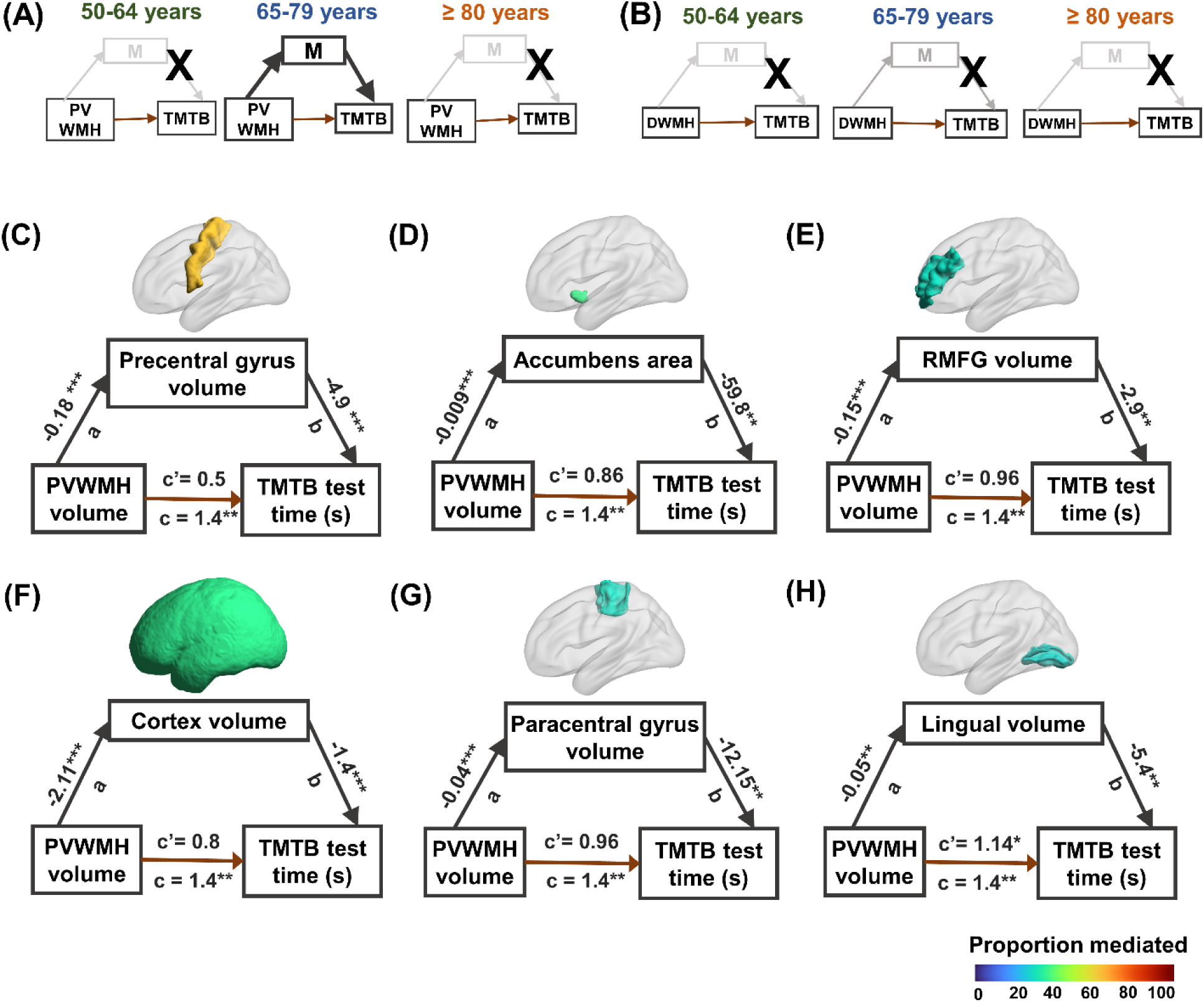
Mediation analysis. (A) Age-group-specific mediation analysis reveals significant brain region mediators in the intermediate age group (65-79 years) with PVWMH as the predictor; no significant mediation was observed in the early and late age groups. **(B)** No significant mediation with DWMH as predictor. **(C-H)** Mediation models show significant mediating brain regions labeled as per the proportion of effect (in %) they mediate in the intermediate age-group (N=271). The values along the paths represent the standardized coefficients: *a* (predictor to mediator), *b* (mediator to outcome), *c’* (direct effect of predictor on the outcome), and *c* (total effect of predictor on the outcome). All paths were adjusted for age. *** p<0.001, ** p<0.01, * p<0.05.

**Table 6.**
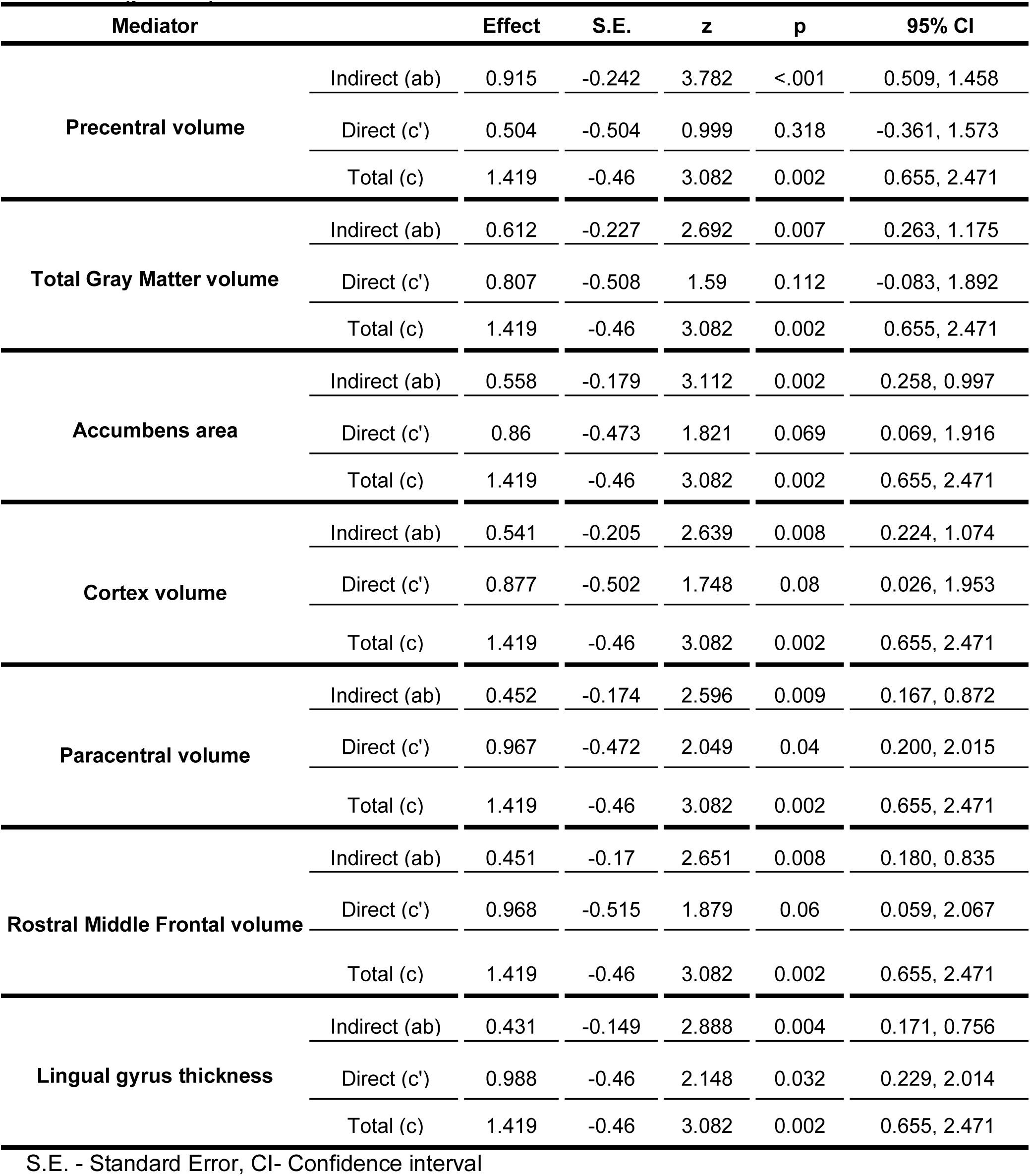
Effect sizes of direct (c’), mediated (ab) and total (c) paths for structural regions as mediator in PVWMH induced TMT-B completion time with significant mediation (p<0.05)

| Mediator |  | Effect | S.E. | z | p | 95% CI |
| --- | --- | --- | --- | --- | --- | --- |
| Precentral volume | Indirect (ab) | 0.915 | -0.242 | 3.782 | <.001 | 0.509, 1.458 |
|  | Direct (c') | 0.504 | -0.504 | 0.999 | 0.318 | -0.361, 1.573 |
|  | Total (c) | 1.419 | -0.46 | 3.082 | 0.002 | 0.655, 2.471 |
| Total Gray Matter volume | Indirect (ab) | 0.612 | -0.227 | 2.692 | 0.007 | 0.263, 1.175 |
|  | Direct (c') | 0.807 | -0.508 | 1.59 | 0.112 | -0.083, 1.892 |
|  | Total (c) | 1.419 | -0.46 | 3.082 | 0.002 | 0.655, 2.471 |
| Accumbens area | Indirect (ab) | 0.558 | -0.179 | 3.112 | 0.002 | 0.258, 0.997 |
|  | Direct (c') | 0.86 | -0.473 | 1.821 | 0.069 | 0.069, 1.916 |
|  | Total (c) | 1.419 | -0.46 | 3.082 | 0.002 | 0.655, 2.471 |
| Cortex volume | Indirect (ab) | 0.541 | -0.205 | 2.639 | 0.008 | 0.224, 1.074 |
|  | Direct (c') | 0.877 | -0.502 | 1.748 | 0.08 | 0.026, 1.953 |
|  | Total (c) | 1.419 | -0.46 | 3.082 | 0.002 | 0.655, 2.471 |
| Paracentral volume | Indirect (ab) | 0.452 | -0.174 | 2.596 | 0.009 | 0.167, 0.872 |
|  | Direct (c') | 0.967 | -0.472 | 2.049 | 0.04 | 0.200, 2.015 |
|  | Total (c) | 1.419 | -0.46 | 3.082 | 0.002 | 0.655, 2.471 |
| Rostral Middle Frontal volume | Indirect (ab) | 0.451 | -0.17 | 2.651 | 0.008 | 0.180, 0.835 |
|  | Direct (c') | 0.968 | -0.515 | 1.879 | 0.06 | 0.059, 2.067 |
|  | Total (c) | 1.419 | -0.46 | 3.082 | 0.002 | 0.655, 2.471 |
| Lingual gyrus thickness | Indirect (ab) | 0.431 | -0.149 | 2.888 | 0.004 | 0.171, 0.756 |
|  | Direct (c') | 0.988 | -0.46 | 2.148 | 0.032 | 0.229, 2.014 |
|  | Total (c) | 1.419 | -0.46 | 3.082 | 0.002 | 0.655, 2.471 |

S.E. - Standard Error, CI- Confidence interval

Moreover, poor performance in TMT-B was also mediated by the precentral gyrus (Effect = 0.915, SE = 0.24, p < 0.001, 95% CI = 0.51, 1.46), paracentral gyrus (Effect = 0.45, SE = 0.17, p = 0.009, 95% CI = 0.17, 0.87) and the nucleus accumbens (Effect = 0.56, SE = 0.18, p = 0.002, 95% CI = 0.26, 0.99) - regions not traditionally linked with TMT-B related executive function. These findings indicate that WMH-induced atrophy in the precentral and paracentral gyrus contributes 64.5% and 31.85% respectively to reaction time delays, a task requiring rapid visual-motor coordination and cognitive flexibility.

Another ANCOVA analysis evaluating the impact of total WMH load on structural volumes in the CN group revealed significant effects on the precentral gyrus (F(3, 384) = 15.1, p < 0.001), nucleus accumbens (F(3, 384) = 18.36, p < 0.001), cerebral cortex (F(3, 384) = 11.36, p < 0.001), paracentral gyrus (F(3, 384) = 9.3, p < 0.001), RMFG (F(3, 384) = 5.96, p < 0.001), and lingual gyrus (F(3, 384) = 4.26, p < 0.001). Comparison of age-adjusted neuroanatomic volumetry measurements across quartiles of PVWMH and DWMH revealed significant structural atrophy. Subjects with >0.9 mL PVWMH (Q2: 0.94 - 2.32 mL) load showed significant reductions in the nucleus accumbens (-8.9%, p=0.009) volume. PVWMH load >2.3 mL i.e., Q3 resulted in significant volume reduction in the precentral (-6.2%, p<0.001), paracentral gyrus (-8.2%, p<0.001), rostral middle frontal gyrus (-8.1%, p<0.001) and cerebral cortex (-6.1%, p<0.001). As PVWMH burden exceeded 6 mL, additional atrophy was detected in the lingual gyrus (-8.6%, p = 0.006) (**Fig. 8**). In contrast, a DWMH load >2.7 mL was associated only with nucleus accumbens atrophy (-12.9%, p = 0.002), with no significant volumetric changes in other brain structures, even at higher DWMH loads.

**Figure 8.**
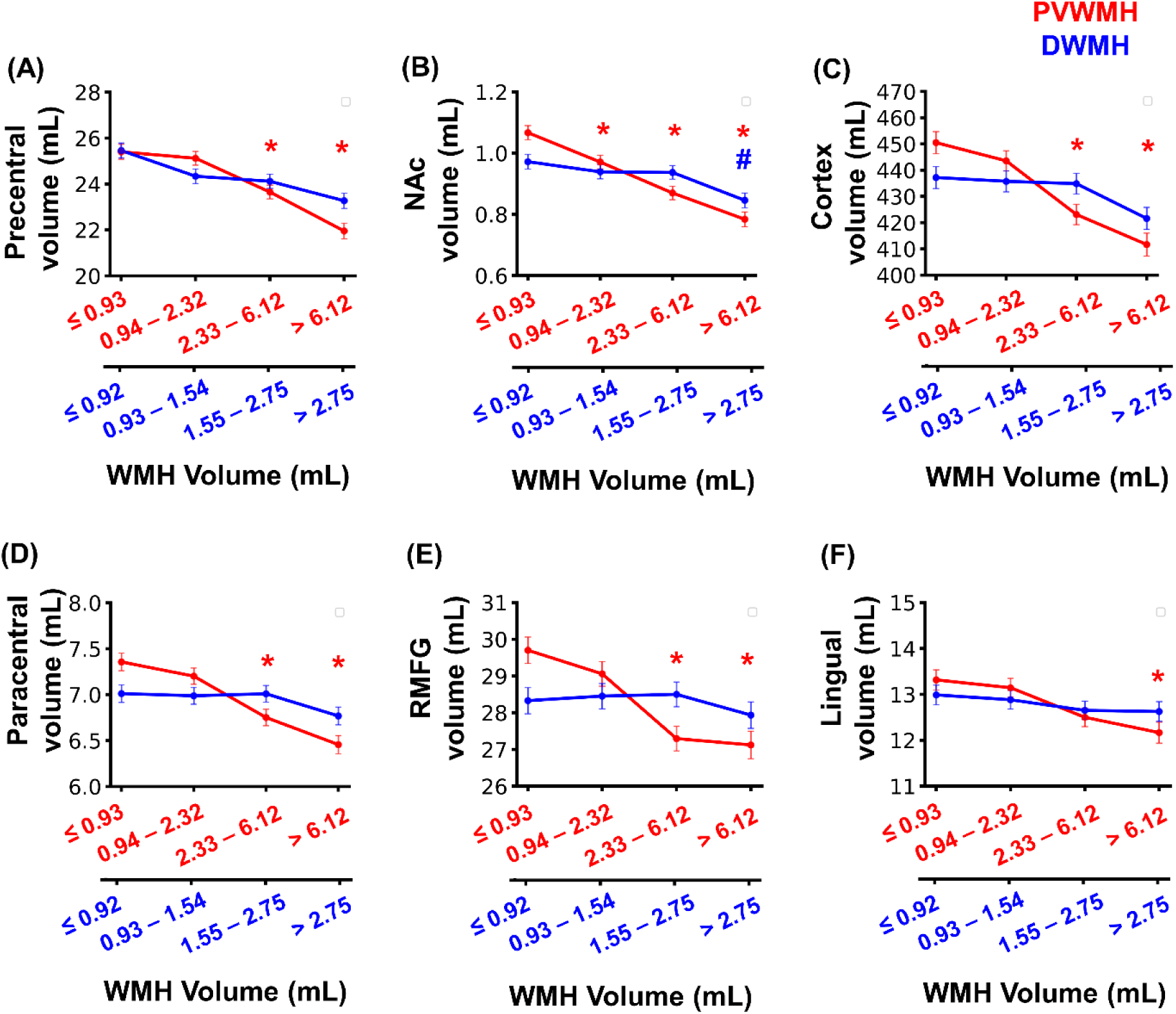
Age-adjusted neuroanatomic volumes of **(A)** Precentral gyrus, **(B)** Nucleus Accumbens, **(C)** Cortex, **(D)** Paracentral gyrus, **(E)** Rostral middle frontal gyrus, and **(F)** Lingual gyrus in CN subjects (N=389) divided into quartiles based on PVWMH volume depicted in red color (0 - 0.93 mL (N=98), 0.94 - 2.32 mL (N=97), 2.33 - 6.12 mL (N=97), and >6.12 mL (N=97)) and DWMH volume (0 - 0.92 mL (N=98), 0.93 - 1.54 mL (N=97), 1.55 - 2.75 mL (N=97), and >2.75 mL (N=97)) depicted in blue color. Neuroanatomic volumes were adjusted for age using ANCOVA, keeping age as covariate, structural volumes as dependent variable and WMH quartiles as fixed factor. Post-hoc pairwise comparisons with Bonferroni correction were applied to identify significant differences between WMH quartiles. The star (*) indicates significant differences compared to the first quartile of PVWMH volume, while the hash (#) denotes significance for DWMH quartiles compared to the first quartile of DWMH. *p < 0.01, #p < 0.01.

Furthermore, longitudinal measurements of WMH and neuroanatomic volumes at baseline and follow-up visits in CN subjects (N = 27) showed that even a small increase in PVWMH load, from none to 1 mL, led to a significant decline in the volume of several brain structures. These included the precentral (p = 0.001), nucleus accumbens (p = 0.002), RMFG (p<0.001), cerebral cortex (p<0.001), paracentral gyri (p<0.001) as well as hippocampus (p = 0.007) and cerebellar white matter (p<0.001) (**Supp.** Fig. 2, **Supp. Table 3**). No significant volume changes were observed in the lingual gyrus, cerebral white matter, or cerebellar cortex.

### Nucleation and growth of WMH lesions

Longitudinal analysis and quantification of segmented WMH masks indicated that WMH lesion progression follows a ‘nucleation and growth model’ (**Fig. 9A**), which may account for the exponential growth of WMH. This model suggests continued nucleation of new lesions along with simultaneous enlargement of existing ones. Additionally, discrete WMH lesions tend to coalesce into larger confluent lesions with age, a pattern especially observed in periventricular WMH (**Fig. 9B**).

**Figure 9.**
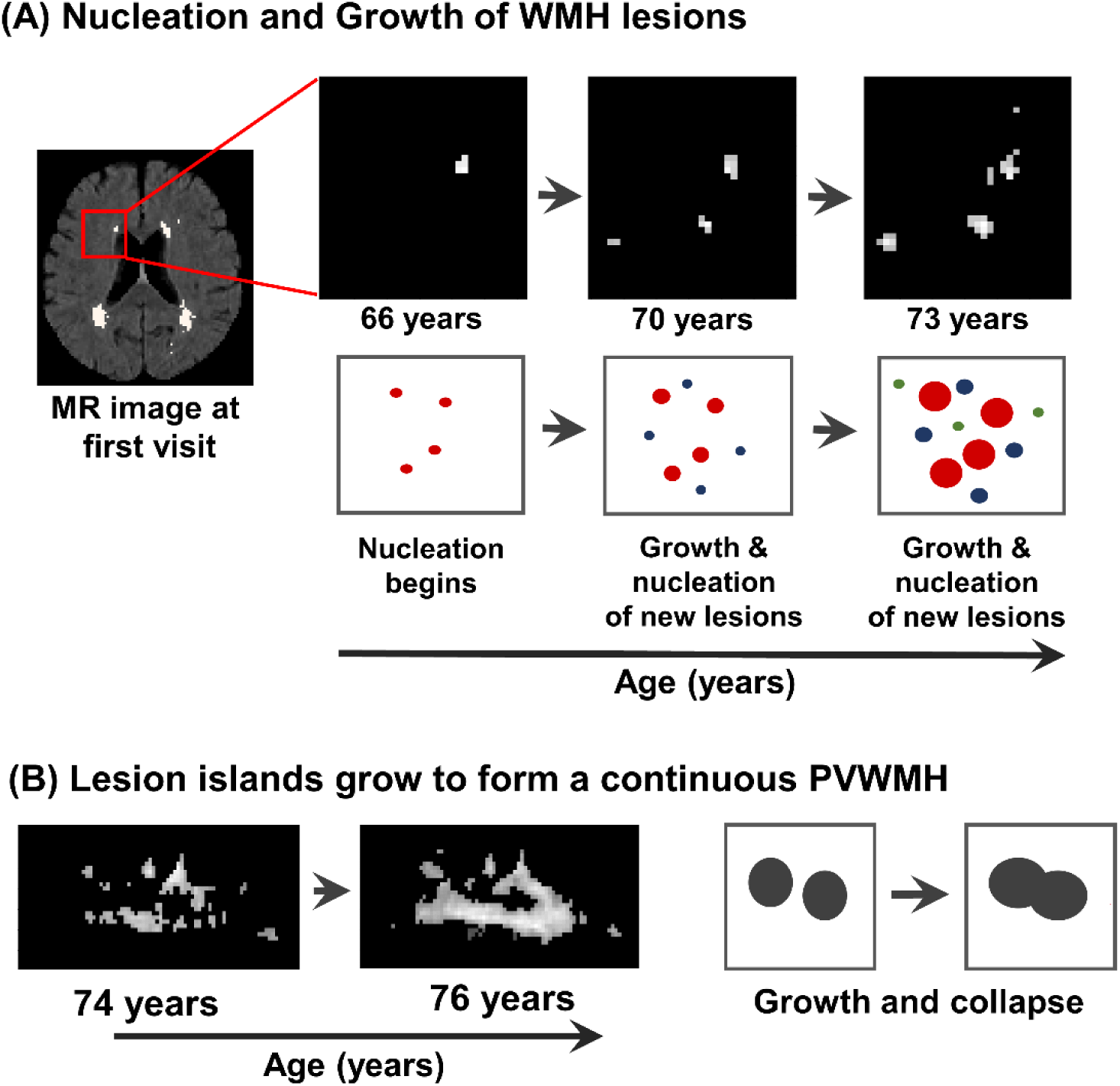
Nucleation and growth of WMH lesions. (A) ROI cutouts from MR images across longitudinal visits showing WMH masks, highlights nucleation of new lesions and the growth of existing lesions with age. **(B)** Discrete WMH lesions in the periventricular region join to form continuous confluent WMH lesions.

## Discussion

It was intriguing to note that despite a significant WMH load in a subset of subjects, the individuals are still classified as cognitively normal (CN). To the best of our knowledge, no previous studies on aging cohorts have explored cognitive and neuroanatomical changes within a cognitive group by segregating subjects based on PVWMH and DWMH volume ranges. In this study, we identify the thresholds of PVWMH and DWMH loads beyond which specific cognitive domains are affected, either directly or through mediation by certain neuroanatomical structures. Our findings suggest that while PVWMH and DWMH loads may not impact global cognition measures, they can affect specific cognitive domains and cause neuroanatomical changes when they exceed a certain threshold for a given age.

Qualitative grading based on Fazekas scores or the shape of WMH (line, halo, or confluent) (Griffanti et al., 2018; Wardlaw et al., 2015) has examined the implications of WMH on tissue microstructure and cognition. However, studies investigating PVWMH and DWMH loads present conflicting findings. For example, van den Heuvel, 2006 suggested that PVWMH at baseline and progression is associated with reduced mental processing speed, but not DWMH load, while Soriano-Raya et al., 2012 reported that DWMH, not PVWMH, is linked to impaired cognitive functions. These contradictory findings likely arise from the qualitative nature of WMH assessments, such as visual ratings, which lack the sensitivity and reliability needed to establish a definitive association between WMH and its impact on cognition and neuroanatomical health.

A few prior quantitative investigations of total WMH have reported a linear increase in WMH with age (Griffanti et al., 2018; van den Heuvel, 2006). However, our findings show that total WMH, PVWMH, and DWMH increase exponentially with age. Interestingly, the progression rate of PVWMH is faster than that of DWMH in CN subjects, although the initial loads of PVWMH and DWMH are similar and remain constant and parallel until around 60 years of age. After the threshold age of approximately 61 years, an inflection point is observed, where the PVWMH increase becomes exponential. A recent study by Kamal et al., 2023 also suggests that WMH accelerates after 60.5 years of age. Similarly, Shen et al., 2024 reported around 60 years as an inflection point in the aging process, marked by significant dysregulation of molecular markers of aging. The convergence of these findings underscores the critical importance of this age threshold as a key juncture in the onset of accelerated vascular and neuroanatomical changes associated with aging. The exponential increase in PVWMH load beyond the age of 61 is likely due to a higher density of confluent PVWMH lesions, which are more punctate in nature before the age of 60. As age progresses, confluent PVWMH lesions increase continuously, along with a further rise in the number of punctate and intermediate lesions. We introduce a “Nucleation and Growth Model” to explain the progression of WMH with age, which illustrates the continuous formation of new lesions alongside the enlargement of existing ones, eventually merging into larger, confluent lesions, particularly in the periventricular region. We propose that individuals with high confluency at an earlier age are more susceptible to the vascular impacts of WMH on cognition and neuroanatomical health.

The mere presence of punctate PVWMH lesions does not necessarily imply a detrimental impact on cognition. However, a PVWMH load >2.3 ml significantly impairs attention and working memory (Digit Span Tests), executive functions (Trail Making Tests), and semantic memory (Animal Naming). At a PVWMH load >6.1 ml, additional impairments become evident in language (Boston naming test), episodic memory (Logical memory -delayed), and global cognition (MMSE), although the changes are minimal for global cognition (-2%) and language domains (-5%). However, we do not observe any significant changes in TMT-A, TMT-B, LMT, BNT and global cognition (MMSE) at any of the DWMH quartiles, while subjects with a DWMH load >2.75 ml show deficits in working memory and semantic memory. Notably, global cognition lacks sensitivity in distinguishing cognitive differences between subjects with and without WMH. A significant decline in working memory and semantic memory is observed when the total WMH (PVWMH + DWMH) load exceeds 4.15 ml. We propose that PVWMH load is an earlier event and it exerts noticeable cognitive debilitations than DWMH load.

The mediation model ascertaining the impact of PVWMH on TMT-B performance shows that WMH-induced atrophy in the RMFG, nucleus accumbens, lingual gyrus, precentral gyrus and paracentral gyrus are significant structural contributors to impaired performance on the TMT-B test. The involvement of RMFG and lingual gyrus in poor TMT-B performance reflects the impact of PVWMH on working memory and visual processing, respectively. Additionally, poor TMT-B performance in subjects with PVWMH is also mediated by atrophy in the precentral and paracentral gyri, which are important for motor coordination, and the nucleus accumbens, which is related to reward processing. The involvement of these motor-related areas likely contributes to coordination delays, while atrophy in the nucleus accumbens may reflect reduced motivation for task completion in subjects with increased PVWMH load. Notably, although PVWMH load is directly and significantly associated with TMT-B performance across all age groups, the mediation effect is significant only in the intermediate age group (65-79 years).

Previous studies have primarily focused on the impact of WMH on cortical thickness in unimpaired subjects (Brugulat-Serrat et al., 2020; Jiménez-Balado et al., 2024; Rizvi et al., 2018) and AD subjects (Zhang et al., 2024). Brugulat-Serrat et al. proposed a mediation model with GM volume as the predictor and WMH volume as the mediator, suggesting that GM volume influences WMH load and thereby reduces executive function. However, we argue that it is the WMH load which beyond a certain threshold is detrimental to structural entities due to increased vascular insult. Indeed, our longitudinal analysis of PVWMH and neuroanatomical volume indicated that subjects with PVWMH progression even to additional 0.5 -1 ml PVWMH load increase at the next visit experienced notable loss in structural volume (in the precentral gyrus, nucleus accumbens, RMFG, paracentral gyrus, cerebral cortex, and hippocampus) compared to subjects with stable or no WMH. The mediation analysis and the neuroanatomic loss associated with WMH progression clearly show that cognitive impairments induced by WMH are manifestations of changes in a specific set of structures related to unique functions. An alternative model with WMH load as the predictor and GM volume as the mediator was not significant, indicating that GM volume alone may not fully account for executive function decline, as observed in poor performance on TMT-B among subjects with PVWMH load exceeding 2.3 ml. Therefore, in light of the reports by Brugulat-Serrat et al., we assert that WMH load exerts cognitive impacts mediated by accelerated neuroanatomic changes in specific domains depending on the WMH load at a given age. DWMH did not show significant neuroanatomic mediation in cognitive losses. Similar to the WMH load thresholds impacting cognitive performance, we identified WMH thresholds associated with structural atrophy. Beyond a PVWMH threshold of >2.3 ml, subjects exhibited significant volumetric reduction in the precentral gyrus, paracentral gyrus, RMFG, and cerebral cortex. At a PVWMH load greater than 0.9 ml, atrophy was evident in the nucleus accumbens, while lingual gyrus atrophy was observed beyond a PVWMH load of 6.1 ml. However, no noticeable changes in structural volumes were observed at any of the DWMH quartiles, except in the nucleus accumbens in subjects with DWMH load exceeding 2.75 ml. In addition to neuroanatomical atrophy and associated cognitive decline, PVWMH is also known to cause significant disruption of long association tracts (Debette et al., 2007).

## Conclusion

PVWMH rises exponentially with age, with a notable inflection point at 61 years. The PVWMH exhibits a faster growth rate than DWMH, particularly in the frontal horn, and accelerates more rapidly in CI and CI-AD groups. A PVWMH burden >2.3 ml is strongly associated with deficits in attention and working memory, executive functioning, and semantic memory. Once the PVWMH burden exceeds 6.1 ml, the cognitive deficits extend to language, episodic memory and global cognition. Significant neuroanatomic atrophy is observed in various brain structures with PVWMH volumes >2.3 ml, whereas DWMH is not significantly associated with neuroanatomic loss. PVWMH deposition precedes DWMH, and an early occurrence of PVWMH load beyond 2.3 ml may be concerning for cognitive health and the future trajectory of an individual’s cognitive function. The ’Nucleation and Growth model’ explains the exponential increase of WMH in cognitively normal subjects. These dynamics need to be evaluated across a large cohort of CI and dementia subjects modeling the trajectory for a very young age to a late life. As vascular risk factors globally are increasing, cerebrovascular burden is expected to rise manifold, henceforth, leading to a much higher cognitive burden than estimates based solely on aging demographics data. This study establishes the importance of quantifying and stratifying PVWMH and DWMH load alongside neuroanatomical volumetry and cognitive scores for a more accurate understanding of aging trajectories and age associated disorders.

## Acknowledgments

The study was funded by Indian Council of Medical Research (ICMR). The MRI and cognitive data were obtained from the National Alzheimer’s Coordinating Centre (NACC) database funded by NIA/NIH Grant U24 AG072122. The NACC database is funded by NIA/NIH Grant U24 AG072122. NACC data are contributed by the NIA-funded ADRCs: P30 AG062429 (PI James Brewer, MD, PhD), P30 AG066468 (PI Oscar Lopez, MD), P30 AG062421 (PI Bradley Hyman, MD, PhD), P30 AG066509 (PI Thomas Grabowski, MD), P30 AG066514 (PI Mary Sano, PhD), P30 AG066530 (PI Helena Chui, MD), P30 AG066507 (PI Marilyn Albert, PhD), P30 AG066444 (PI John Morris, MD), P30 AG066518 (PI Jeffrey Kaye, MD), P30 AG066512 (PI Thomas Wisniewski, MD), P30 AG066462 (PI Scott Small, MD), P30 AG072979 (PI David Wolk, MD), P30 AG072972 (PI Charles DeCarli, MD), P30 AG072976 (PI Andrew Saykin, PsyD), P30 AG072975 (PI David Bennett, MD), P30 AG072978 (PI Neil Kowall, MD), P30 AG072977 (PI Robert Vassar, PhD), P30 AG066519 (PI Frank LaFerla, PhD), P30 AG062677 (PI Ronald Petersen, MD, PhD), P30 AG079280 (PI Eric Reiman, MD), P30 AG062422 (PI Gil Rabinovici, MD), P30 AG066511 (PI Allan Levey, MD, PhD), P30 AG072946 (PI Linda Van Eldik, PhD), P30 AG062715 (PI Sanjay Asthana, MD, FRCP), P30 AG072973 (PI Russell Swerdlow, MD), P30 AG066506 (PI Todd Golde, MD, PhD), P30 AG066508 (PI Stephen Strittmatter, MD, PhD), P30 AG066515 (PI Victor Henderson, MD, MS), P30 AG072947 (PI Suzanne Craft, PhD), P30 AG072931 (PI Henry Paulson, MD, PhD), P30 AG066546 (PI Sudha Seshadri, MD), P20 AG068024 (PI Erik Roberson, MD, PhD), P20 AG068053 (PI Justin Miller, PhD), P20 AG068077 (PI Gary Rosenberg, MD), P20 AG068082 (PI Angela Jefferson, (which was not certified by peer review) is the author/funder. All rights reserved. No reuse allowed without permission. bioRxiv preprint doi: https://doi.org/10.1101/2023.09.05.556337; this version posted September 5, 2023. The copyright holder for this preprint PhD), P30 AG072958 (PI Heather Whitson, MD), P30 AG072959 (PI James Leverenz, MD).

