## Supplementary data for "Periventricular and Deep White Matter Hyperintensity Thresholds in Aging: Exponential Progression, Cognitive Decline, and Neuroanatomic Atrophy"

**Supplementary Figure 1.**

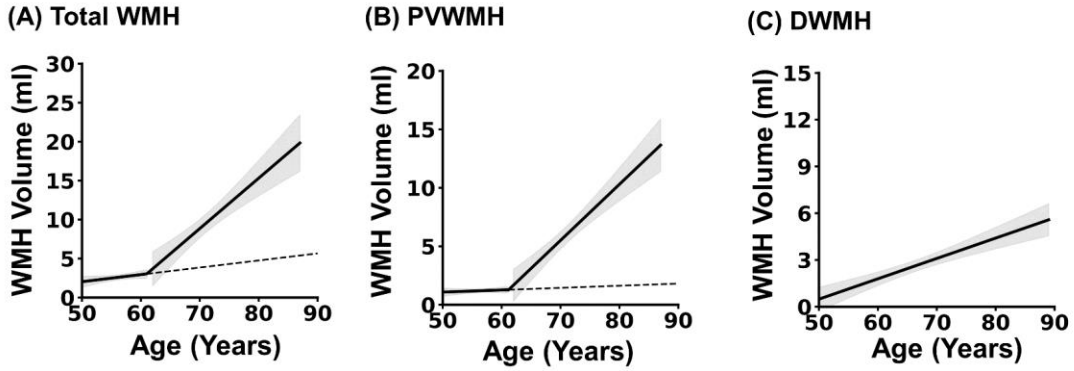


**Supplementary Figure 1.** Identification of the inflection point using a piecewise linear regression algorithm, followed by linear regression analysis before and after the inflection point in CN subjects for **(A)** total WMH, **(B)** PVWMH volume with age. No inflection point was observed for **(C)** DWMH volume with age. The shaded gray area represents a 95% confidence interval. The dashed line represents the pre-inflection path, while the solid line denotes the linear slope after the inflection point.

### Supplementary Figure 2

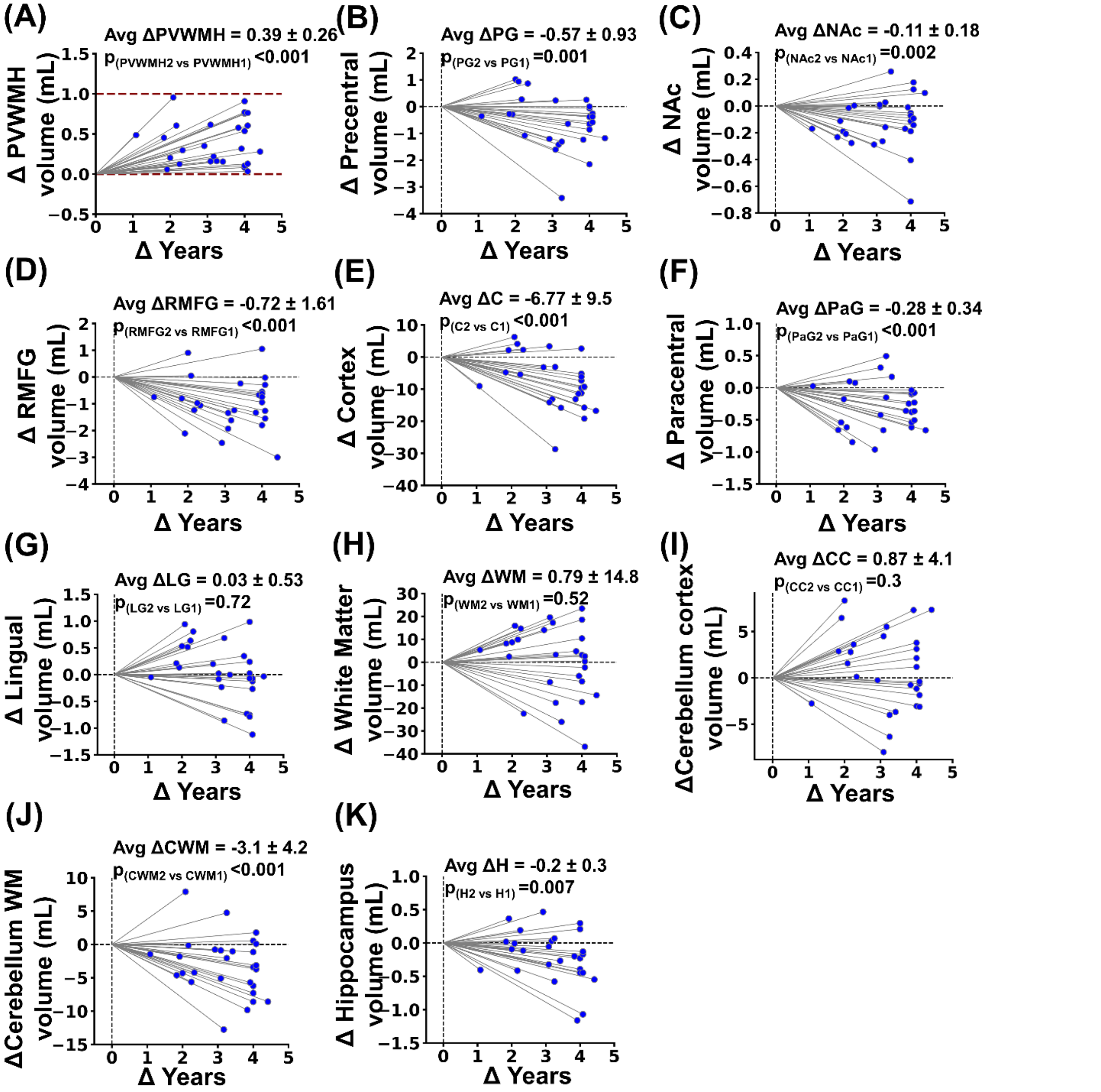


**Supplementary Figure 2. Longitudinal structural volume changes in subjects (N=27) with a 0–1 mL increase in ΔPVWMH volume. (A)** ΔPVWMH volume shows an average increase of 0.39 ± 0.26 mL, with red dashed lines indicating the 0–1 mL ΔPVWMH range. **(B-F, J-K)** Significant decrease in volume were observed in the Precentral Gyrus (PG), Nucleus Accumbens (NAc), Rostral Middle Frontal Gyrus (RMFG), Cortex (C), Paracentral Gyrus (PaG), Cerebellar white matter and Hippocampal volume between the visits. No significant changes were noted in **(G)** Lingual, **(H)** Cerebral White Matter and **(I)** cerebellar cortex volumes. The x-axes represent the interval (years) between visits. Statistical significance between visits was assessed using the Wilcoxon rank-sum test. Values are presented as the mean ± standard deviation of the change in structural volume at the follow-up visit.

### Supplementary Table 1: Instantaneous rates of PVWMH and DWMH with Age

| Age | CN | | | CI | | | CI-AD | | |
|---|---|---|---|---|---|---|---|---|---|
| | PVWMH rate | DWMH rate | PV / Deep | PVWMH rate | DWMH rate | PV / Deep | PVWMH rate | DWMH rate | PV / Deep |
| **50** | 0.087 | 0.054 | **1.625** | 0.157 | 0.083 | **1.888** | 0.144 | 0.086 | **1.667** |
| **51** | 0.093 | 0.056 | **1.659** | 0.164 | 0.087 | **1.883** | 0.150 | 0.090 | **1.666** |
| **52** | 0.099 | 0.059 | **1.694** | 0.171 | 0.091 | **1.879** | 0.157 | 0.094 | **1.666** |
| **53** | 0.106 | 0.061 | **1.730** | 0.179 | 0.095 | **1.874** | 0.164 | 0.099 | **1.665** |
| **54** | 0.114 | 0.064 | **1.767** | 0.187 | 0.100 | **1.870** | 0.172 | 0.103 | **1.665** |
| **55** | 0.122 | 0.067 | **1.804** | 0.195 | 0.104 | **1.865** | 0.180 | 0.108 | **1.664** |
| **56** | 0.130 | 0.071 | **1.842** | 0.203 | 0.109 | **1.861** | 0.188 | 0.113 | **1.664** |
| **57** | 0.139 | 0.074 | **1.881** | 0.212 | 0.114 | **1.856** | 0.197 | 0.119 | **1.663** |
| **58** | 0.149 | 0.077 | **1.921** | 0.221 | 0.119 | **1.852** | 0.206 | 0.124 | **1.663** |
| **59** | 0.159 | 0.081 | **1.961** | 0.231 | 0.125 | **1.848** | 0.216 | 0.130 | **1.662** |
| **60** | 0.170 | 0.085 | **2.003** | 0.241 | 0.131 | **1.843** | 0.226 | 0.136 | **1.662** |
| **61** | 0.182 | 0.089 | **2.045** | 0.252 | 0.137 | **1.839** | 0.236 | 0.142 | **1.661** |
| **62** | 0.194 | 0.093 | **2.088** | 0.263 | 0.143 | **1.834** | 0.247 | 0.149 | **1.661** |
| **63** | 0.208 | 0.097 | **2.132** | 0.274 | 0.150 | **1.830** | 0.259 | 0.156 | **1.660** |
| **64** | 0.222 | 0.102 | **2.177** | 0.286 | 0.157 | **1.826** | 0.271 | 0.163 | **1.660** |
| **65** | 0.238 | 0.107 | **2.223** | 0.298 | 0.164 | **1.821** | 0.283 | 0.171 | **1.659** |
| **66** | 0.254 | 0.112 | **2.270** | 0.311 | 0.171 | **1.817** | 0.296 | 0.179 | **1.659** |
| **67** | 0.272 | 0.117 | **2.318** | 0.325 | 0.179 | **1.812** | 0.310 | 0.187 | **1.658** |
| **68** | 0.290 | 0.123 | **2.367** | 0.339 | 0.188 | **1.808** | 0.324 | 0.196 | **1.658** |
| **69** | 0.311 | 0.129 | **2.417** | 0.354 | 0.196 | **1.804** | 0.339 | 0.205 | **1.657** |
| **70** | 0.332 | 0.135 | **2.468** | 0.369 | 0.205 | **1.799** | 0.355 | 0.214 | **1.657** |
| **71** | 0.355 | 0.141 | **2.520** | 0.386 | 0.215 | **1.795** | 0.372 | 0.224 | **1.656** |
| **72** | 0.380 | 0.148 | **2.573** | 0.402 | 0.225 | **1.791** | 0.389 | 0.235 | **1.656** |
| **73** | 0.406 | 0.155 | **2.628** | 0.420 | 0.235 | **1.787** | 0.407 | 0.246 | **1.655** |
| **74** | 0.434 | 0.162 | **2.683** | 0.438 | 0.246 | **1.782** | 0.426 | 0.257 | **1.655** |
| **75** | 0.464 | 0.169 | **2.740** | 0.457 | 0.257 | **1.778** | 0.446 | 0.269 | **1.654** |
| **76** | 0.496 | 0.177 | **2.798** | 0.477 | 0.269 | **1.774** | 0.466 | 0.282 | **1.654** |
| **77** | 0.531 | 0.186 | **2.857** | 0.498 | 0.282 | **1.769** | 0.488 | 0.295 | **1.653** |
| **78** | 0.568 | 0.195 | **2.917** | 0.520 | 0.294 | **1.765** | 0.510 | 0.309 | **1.653** |
| **79** | 0.607 | 0.204 | **2.979** | 0.543 | 0.308 | **1.761** | 0.534 | 0.323 | **1.652** |
| **80** | 0.649 | 0.213 | **3.042** | 0.566 | 0.322 | **1.757** | 0.559 | 0.338 | **1.652** |
| **81** | 0.694 | 0.223 | **3.106** | 0.591 | 0.337 | **1.753** | 0.585 | 0.354 | **1.651** |
| **82** | 0.742 | 0.234 | **3.172** | 0.617 | 0.353 | **1.748** | 0.612 | 0.371 | **1.651** |
| **83** | 0.794 | 0.245 | **3.239** | 0.644 | 0.369 | **1.744** | 0.640 | 0.388 | **1.650** |
| **84** | 0.849 | 0.257 | **3.307** | 0.672 | 0.386 | **1.740** | 0.670 | 0.406 | **1.650** |
| **85** | 0.907 | 0.269 | **3.377** | 0.701 | 0.404 | **1.736** | 0.701 | 0.425 | **1.649** |

| | | | | | | | | | |
|---|---|---|---|---|---|---|---|---|---|
| **86** | 0.970 | 0.281 | **3.448** | 0.732 | 0.422 | **1.732** | 0.733 | 0.445 | **1.649** |
| **87** | 1.037 | 0.295 | **3.521** | 0.763 | 0.442 | **1.728** | 0.767 | 0.466 | **1.648** |
| **88** | 1.109 | 0.309 | **3.595** | 0.797 | 0.462 | **1.723** | 0.803 | 0.487 | **1.648** |
| **89** | 1.186 | 0.323 | **3.671** | 0.831 | 0.484 | **1.719** | 0.840 | 0.510 | **1.647** |
| **90** | 1.268 | 0.338 | **3.749** | 0.868 | 0.506 | **1.715** | 0.879 | 0.534 | **1.647** |
| **91** | 1.356 | 0.354 | **3.828** | 0.906 | 0.529 | **1.711** | 0.920 | 0.559 | **1.646** |
| **92** | 1.450 | 0.371 | **3.909** | 0.945 | 0.554 | **1.707** | 0.962 | 0.585 | **1.646** |
| **93** | 1.551 | 0.389 | **3.991** | 0.986 | 0.579 | **1.703** | 1.007 | 0.612 | **1.645** |
| **94** | 1.658 | 0.407 | **4.076** | 1.029 | 0.606 | **1.699** | 1.054 | 0.641 | **1.645** |
| **95** | 1.773 | 0.426 | **4.162** | 1.074 | 0.634 | **1.695** | 1.102 | 0.670 | **1.644** |
| **Mean ± SD** | 0.57 ±0.5 | 0.18 ±0.1 | 2.7 ±0.8 | 0.48 ±0.3 | 0.27 ±0.2 | 1.79 ±0.1 | 0.47 ±0.3 | 0.29 ±0.2 | 1.66 ±0.01 |

SD- Standard deviation

### Supplementary Table 2. Dominance analysis results

| Neuropsychological test scores (Dependent variable) | Regional WMH Volume (Independent variable) | Interactional Dominance | Individual Dominance | Average Partial Dominance | Total Dominance | Percentage Relative Importance |
|---|---|---|---|---|---|---|
| MMSE | PVWMH | 0.044 | 0.064 | 0.051 | 0.052 | 60.821 |
| | Frontal WMH | 0.001 | 0.019 | 0.005 | 0.007 | 8.241 |
| | Temporal WMH | 0.011 | 0.001 | 0.007 | 0.007 | 7.804 |
| | Parietal WMH | 0.002 | 0.034 | 0.014 | 0.014 | 18.189 |
| | Occipital WMH | 0.007 | 0.001 | 0.004 | 0.004 | 4.944 |
| Boston Naming Test | PVWMH | 0.035 | 0.035 | 0.034 | 0.035 | 72.086 |
| | Frontal WMH | 0.000 | 0.010 | 0.004 | 0.004 | 8.952 |
| | Temporal WMH | 0.007 | 0.002 | 0.004 | 0.004 | 8.689 |
| | Parietal WMH | 0.005 | 0.000 | 0.003 | 0.003 | 5.648 |
| | Occipital WMH | 0.000 | 0.006 | 0.002 | 0.002 | 4.625 |
| Logical Memory Test | PVWMH | 0.028 | 0.049 | 0.035 | 0.037 | 50.223 |
| | Frontal WMH | 0.000 | 0.024 | 0.007 | 0.009 | 12.115 |
| | Temporal WMH | 0.000 | 0.010 | 0.002 | 0.003 | 4.590 |
| | Parietal WMH | 0.000 | 0.022 | 0.008 | 0.009 | 12.366 |
| | Occipital WMH | 0.022 | 0.002 | 0.017 | 0.015 | 20.705 |
| Animal Naming Test | PVWMH | 0.061 | 0.097 | 0.069 | 0.073 | 67.848 |
| | Frontal WMH | 0.009 | 0.013 | 0.006 | 0.008 | 7.252 |
| | Temporal WMH | 0.001 | 0.029 | 0.007 | 0.010 | 9.226 |
| | Parietal WMH | 0.000 | 0.034 | 0.008 | 0.012 | 11.059 |
| | Occipital WMH | 0.000 | 0.017 | 0.003 | 0.005 | 4.616 |
| Digit Span Test - Forward | PVWMH | 0.025 | 0.058 | 0.034 | 0.037 | 56.844 |
| | Frontal WMH | 0.005 | 0.009 | 0.003 | 0.005 | 7.565 |

| | | | | | | |
|---|---|---|---|---|---|---|
| | Temporal WMH | 0.000 | 0.014 | 0.002 | 0.004 | 5.996 |
| | Parietal WMH | 0.003 | 0.037 | 0.012 | 0.016 | 23.679 |
| | Occipital WMH | 0.000 | 0.013 | 0.002 | 0.004 | 5.916 |
| Digit Span Test - Backward | PVWMH | 0.032 | 0.088 | 0.047 | 0.052 | 55.285 |
| | Frontal WMH | 0.001 | 0.034 | 0.007 | 0.011 | 11.708 |
| | Temporal WMH | 0.005 | 0.011 | 0.003 | 0.005 | 4.882 |
| | Parietal WMH | 0.001 | 0.054 | 0.015 | 0.020 | 21.108 |
| | Occipital WMH | 0.001 | 0.021 | 0.004 | 0.007 | 7.018 |
| Trail Making Test - A | PVWMH | 0.073 | 0.077 | 0.071 | 0.073 | 72.609 |
| | Frontal WMH | 0.008 | 0.007 | 0.006 | 0.006 | 6.479 |
| | Temporal WMH | 0.002 | 0.025 | 0.008 | 0.010 | 10.249 |
| | Parietal WMH | 0.008 | 0.011 | 0.007 | 0.008 | 8.186 |
| | Occipital WMH | 0.001 | 0.007 | 0.001 | 0.002 | 2.477 |
| Trail Making Test - B | PVWMH | 0.087 | 0.133 | 0.097 | 0.102 | 72.365 |
| | Frontal WMH | 0.004 | 0.026 | 0.006 | 0.010 | 6.967 |
| | Temporal WMH | 0.002 | 0.041 | 0.010 | 0.015 | 10.279 |
| | Parietal WMH | 0.000 | 0.027 | 0.004 | 0.008 | 5.759 |
| | Occipital WMH | 0.000 | 0.022 | 0.004 | 0.007 | 4.631 |

**Supplementary Table 3: Longitudinal change in structural volume in subjects with 0 to 1mL increase in Δ PVWMH volume**

| Structural volume | Avg volume At Visit 1 | Avg volume At Visit 2 | Avg Δ Change in volume | $p_{(V1\ vs\ V2)}$ |
|---|---|---|---|---|
| PVWMH | 0.68 ± 0.53 | 1.08 ± 0.62 | 0.40 ± 0.27 | **7.45e-09** |
| DWMH | 0.87 ± 0.51 | 1.46 ± 2.12 | 0.60 ± 2.08 | 0.79 |
| Precentral | 26.46 ± 2.88 | 25.89 ± 2.95 | -0.58 ± 0.93 | **0.001** |
| Accumbens | 1.16 ± 0.24 | 1.04 ± 0.15 | -0.11 ± 0.19 | **0.002** |
| RMFG | 31.26 ± 5.65 | 30.54 ± 5.79 | -0.72 ± 1.62 | **0.0003** |
| Total Gray Matter | 623.18 ± 58.68 | 616.40 ± 60.38 | -6.77 ± 9.54 | **0.0005** |
| Paracentral | 7.63 ± 0.91 | 7.34 ± 0.78 | -0.29 ± 0.35 | **0.0002** |
| Hippocampus | 8.11 ± 0.67 | 7.90 ± 0.83 | -0.21 ± 0.37 | **0.0076** |
| Lingual gyrus | 13.72 ± 2.07 | 13.76 ± 2.38 | 0.04 ± 0.54 | 0.073 |
| Cerebral WM | 447.04 ± 57.15 | 447.84 ± 63.72 | 0.80 ± 14.89 | 0.052 |
| Cerebellum Cortex | 102.63 ± 9.82 | 103.51 ± 10.99 | 0.88 ± 4.12 | 0.3 |
| Cerebellum WM | 33.09 ± 4.29 | 29.97 ± 5.66 | -3.12 ± 4.27 | **0.003** |
